# Neural relational inference to learn allosteric long-range interactions in proteins from molecular dynamics simulations

**DOI:** 10.1101/2021.01.20.427459

**Authors:** Jingxuan Zhu, Juexin Wang, Weiwei Han, Dong Xu

**Affiliations:** Key Laboratory for Molecular Enzymology and Engineering of Ministry of Education, School of Life Science, Jilin University, Changchun, China; Department of Electrical Engineering and Computer Science, Bond Life Sciences Center, University of Missouri, Columbia, MO, United States

## Abstract

Protein allostery is a biological process facilitated by spatially long-range intra-protein communication, whereby ligand binding or amino acid mutation at a distant site affects the active site remotely. Molecular dynamics (MD) simulation provides a powerful computational approach to probe the allostery effect. However, current MD simulations cannot reach the time scales of whole allostery processes. The advent of deep learning made it possible to evaluate both spatially short and long-range communications for understanding allostery. For this purpose, we applied a neural relational inference (NRI) model based on a graph neural network (GNN), which adopts an encoder-decoder architecture to simultaneously infer latent interactions to probe protein allosteric processes as dynamic networks of interacting residues. From the MD trajectories, this model successfully learned the long-range interactions and pathways that can mediate the allosteric communications between the two distant binding sites in the Pin1, SOD1, and MEK1 systems.

## Introduction

Many protein functions are regulated by specific dynamic biomolecular processes, such as allostery, protein folding/unfolding, and protein activation. A biomolecular motion can be considered as a dynamic system driven by atomic/residue interactions. Molecular dynamics (MD) simulations can directly probe the biomolecular motion but may fail to capture meaningful functional information due to the limited time scale of simulation, as well as the high dimensionality and complexity of 3D trajectory data. To extract the biological information from the massive data, statistical techniques such as principal component analysis (PCA)^1^ and cross-correlation analysis^2^ have been applied in MD analyses. PCA reduces the data dimension while maintaining essential information, making the biomolecular motion more interpretable and allowing for visualization. Cross-correlation analysis assesses the extent to which the atomic/residue fluctuations are correlated with one another by examining the magnitude of all pairwise cross-correlation coefficients. However, these methods are inherently restricted to the linear relationships between data features^2,3^, and therefore miss the nonlinear correlation in dynamics closely related to long-range communication in protein^4^. As a result, many challenging MD analysis problems lack good methods to probe long-range communications and nonlinear effects. For example, allosteric communication is ubiquitous in proteins, but how signals are transmitted over long distances within a protein or across different protein molecules has been a conundrum for a long time^5^.

To model protein allosteric communication, many graph models have been developed. Generally, a protein can be mapped to a graph, in which each node represents a residue and each weighted edge represents an interaction between two nodes. The shortest paths between the allosteric site and active sites in protein may be important for propagating signal in the allosteric communication. Earlier graph models used a static crystal structure to calculate the shortest path lengths between one residue to other residues^6^. Later, the information from MD simulations was used to define the interaction graph and shortest path^7^ but not sufficient to characterize allostery.

The advent of deep learning provided new opportunities to explore allosteric effects. The emerging graph neural networks (GNN) is designed to model data systems in graphs and it achieves great success in many graph-related problems^8–10^. Recently, GNN has been proved its great promise in modeling complex dynamic systems in traffic scenes, dynamic physical systems, and computer vision tasks using implicit interaction models with message passing^11–13^ or attention mechanisms^14^. More notably, an unsupervised neural relational inference (NRI) model can infer an explicit interaction structure while simultaneously predicting the dynamic model in physical simulations^15^. This model trains a form of a variational autoencoder using the motion capture data to model dynamics of the input system, in which the learned embedding (latent code) translates the underlying interaction into an interpretable graph structure and predicts time-related dynamics.

We adapted the NRI model (Fig. 1) to understand the allosteric pathway mediating remote regulation from ligand binding or mutation site to the active center in a protein. Based on the trajectories running from MD simulations, we formulated the protein allosteric processes as the dynamic networks of interacting residues. This model uses GNN to learn the embedding of the network dynamics by minimizing the reconstruction error between the reconstructed and input trajectories, then infers edges between residues represented by latent variables. The learned embedding inherently abstracts the essential roles of the key residues in the conformational transition, which helps decipher the mechanism of protein allostery. In this study, we performed MD simulations for three allosteric systems, i.e., (*i*) the allosteric regulation of Pin1 induced by ligand binding, (*ii*) the conformational transition of SOD1 by G93A amyotrophic lateral sclerosis-linked mutation, and (*iii*) the activation of MEK1 by oncogenic mutations. Compared to the conventional MD analysis, our results show that the GNN-based model can learn interpretable interaction patterns and paths in a nonlinear protein system. To the best of our knowledge, this study is the first attempt to use GNN, particularly NRI to analyze MD simulations in biological systems.

**Fig. 1.**
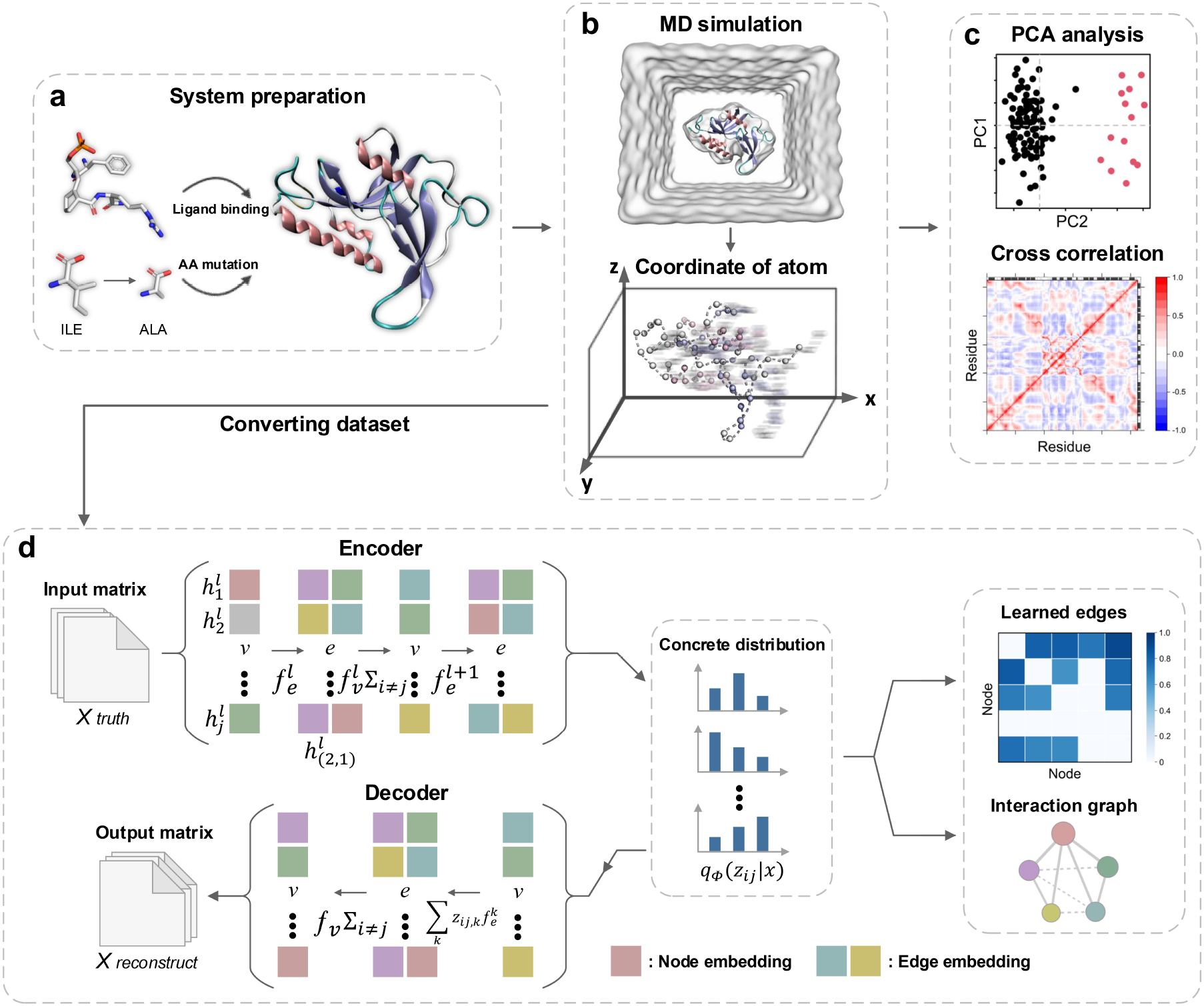
The process of inferring an interaction graph by reconstructing an MD simulation trajectory. **a**, The system preparation of a ligand-binding complex or mutant protein structure with allostery. **b**, The MD simulation of prepared allosteric system to obtain the trajectory with the dynamic 3D coordinates. **c**, The conventional analysis for the trajectory, such as PCA or cross-correlation calculation. **d**, The architecture of NRI model, which consists of two jointly trained components, i.e., an encoder infers a factorized distribution *q*_*ϕ*_(z|x) over the latent interactions based on input trajectories, and a decoder reconstructs several time steps of the dynamic systems given the latent graph learned from the encoder. Based on the MD trajectory, the NRI model formulates the protein allosteric process as a dynamic network of interacting residues. The interaction graph learned from this model is compared to the conventional analysis for a better understanding of the allosteric pathway in the protein.

## Results

### Pathways mediate inter-domain allosteric communication in Pin1

Pin1 utilizes allostery to alter functional activities by changing the local effective modulus of protein without conformational changes^16^. Pin1 as an attractive therapeutic target, contains an inactive N-terminal Trp-Trp (WW) domain (residues 1-39) and an enzymatically active C-terminal peptidyl-prolyl isomerase (PPIase) domain (residues 50-163) connected by a linker (residues 40-49)^17^. The PPIase domain is composed of a PPIase core (α4-helix and β4-β7 strands), α1-α3 helices, and a catalytic loop (Fig. 2a). While both domains bind phospho-Ser/Thr-Pro containing substrate motifs, only the PPIase domain can isomerize peptidyl-prolyl bond through the catalytic site to control post-phosphorylation in regulating Pin1 function^18^. Moreover, the substrate-binding affinities of full-length Pin1 are generally higher than the isolated PPIase domain, suggesting that the non-catalytic WW domain has the potential to remotely modulate the catalytic activity of the PPIase domain^19^. We performed two simulations of Pin1 in apo and FFpSPR-bound form to evaluate the long-range effect of substrate binding to the WW-domain on the flexibility of the protein backbone. The root-mean-square deviation (RMSD) and root-mean-square fluctuation (RMSF) values for the simulations (Fig. 2a and Fig. S1a) show that the apo form exhibits high flexibility in the WW domain (β1-β2), catalytic loop, α2-helix, and the PPIase core (β5/α4). In contrast, the flexibilities of these domains are significantly quenched when the FFpSPR binds to the WW domain, indicating that the ligand binding to the WW domain not only stabilizes the conformation of the WW domain but also remotely regulates other domains to make them more rigid.

**Fig. 2.**
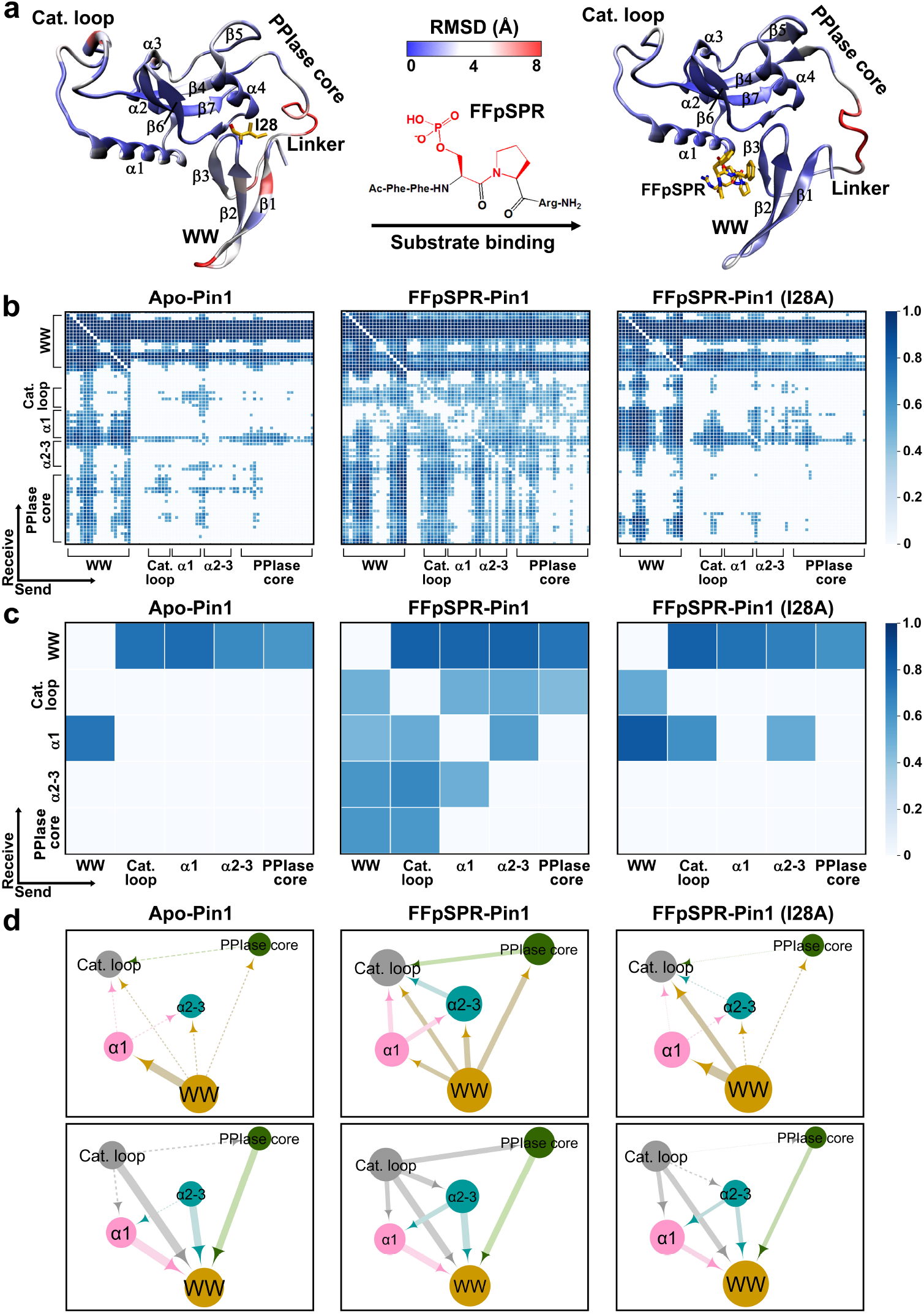
Change of domain communications upon binding and mutation in Pin1. **a**, The flexibility of the protein backbone of apo-Pin1 (*left*) and FFpSPR-Pin1 (*right*), where the color scale represents the backbone RMSD. **b**, Distribution of learned edges between residues in the MD simulations of apo-Pin1 (*left*), FFpSPR-Pin1 (*middle*), and I28A FFpSPR-Pin1 (*right*). **c**, Distribution of learned edges between domains/blocks in the MD simulations of apo-Pin1 (*left*), FFpSPR-Pin1 (*middle*), and I28A FFpSPR-Pin1 (*right*). **d**, The interacted domains/blocks mapped from the learned edges. The size of a node represents the number of edges that directly connect to the node. The thickness of an edge represents the strength of the interaction. The arrowhead denotes the directionality of a learned edge, i.e. the influence from the one starting domain to the ending domain.

To explore the pathways mediating the allosteric regulation of the PPIase domain by its WW domain, we introduced a dynamic model of allostery learned by the NRI model. We utilized the MD trajectories to generate training, validation, and test samples, and trained the NRI model with an encoder and decoder (See Methods for details). We selected 50 uniform timesteps in the 4 ns interval as the ground truth to the model and reconstructed these timesteps. The ground truth and reconstructed trajectories (Supplementary Video 1), the RMSF values (Fig. S2), and the MSE values (Table S1) show that the model can correctly reconstruct the trajectory of motion with small errors while learning the interaction graph.

According to the distribution of learned edges between residues (Fig. 2b), we integrated adjacent residues as blocks for a more straightforward observation of the interactions (Fig. 2c and d). It can be seen that the learned edges often occur between the WW domain and other domains, suggesting that the WW domain is the key element in protein movement. According to these learned edges, we calculated the shortest pathways from residues in the WW domain to residues in the catalytic loop (Table S2). Notably, when the FFpSPR binds to the WW domain, the correlation between the WW domain and PPIase core is reinforced to launch the first two types of pathways, i.e., from the WW domain to Q131 or P133 in the PPIase core (Fig. 3a, *left and middle*). Then the direct coupling between the PPIase core and catalytic loop completes the allosteric communication from the WW domain to the catalytic loop via the WW-PPIase core link (Fig. 3a, *left and middle*). Besides, the FFpSPR binding strengthens another communication from the WW domain via K97 in the α1-helix and S105/C113 in the α2-3 helices to the catalytic loop (Fig. 3a, *right*). In these pathways, the Q131 and P133 in the PPIase core make greater effects on signal propagating than the residues in α-helices. Furthermore, the MD trajectories of different time scales for the FFpSPR-bound Pin1 (Fig. S3) indicate that no allosteric communication is found at first, and the α-helices and PPIase core strengthen the formation of the pathway as the simulation time increased. Finally, the long-range interactions are formed to tighten the catalytic loop of Pin1. However, in the absence of ligand binding, no pathway is found from the WW domain to the catalytic loop. Although the WW domain can interact with the α1-helix, the communication cannot pass from the α1-helix to the catalytic loop (Fig. 3b and Table S2).

**Fig. 3.**
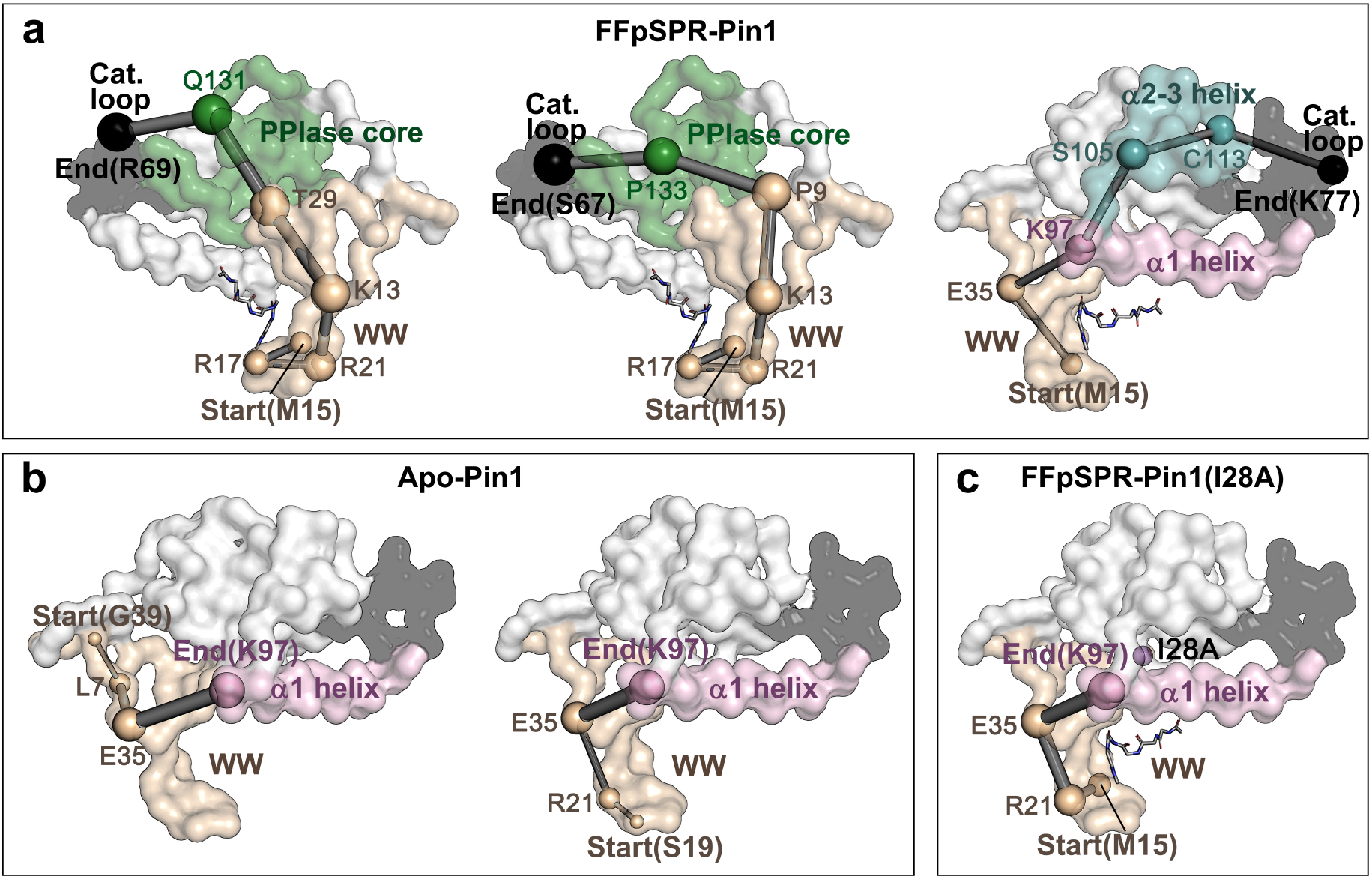
Pathways mediate inter-domain allosteric communications in Pin1, obtained from shortest pathway calculation. **a**, The allosteric pathways mediating remote communication from WW-domain (WW) to catalytic loop in FFpSPR-Pin1: the *left* is from the WW through Q131 in PPIase core to R69 in the catalytic loop; the *middle* is from the WW through P133 in the PPIase core to S67 in the catalytic loop; the *right* is from the WW through K97 in α1-helix and S105/C113 in α2-3 helices to K77 in the catalytic loop. We used the residues in the WW domain as the starting point and residues in the catalytic loop as the ending points to present the shortest pathways (More pathways are shown in Table S2). **b**, The two pathways in apo-Pin1: the *left* is from G39 through L7/E35 to K97 in the α1-helix; the *right* is from S19 through R21/E35 to K97 in the α1-helix. **c**, The pathway in I28A FFpSPR-Pin1: from M15 through R21/E35 to K97 in the α1-helix. The size of a node represents the number of edges that directly connect to the node. The thickness of an edge represents the strength of the interaction.

An NMR study^20^ shows that the I28A mutation can weaken inter-domain interactions, responsible for enhancing the catalytic loop’s flexibility to reduce the substrate affinity for the catalytic site. Since the I28A mutation locates at the inter-domain interface (i.e., the top of the β2-β3 sheet), this mutation may affect the inter-domain coupling between the WW domain and the PPIase domain. We simulated the I28A Pin1 of the FFpSPR-bound form and the trajectory’s RMSF value shows that the I28A mutation increases the mobility of the whole protein structure, especially in the WW domain, catalytic loop, and the α1-α3 helices (Fig. S1b). The learned interaction graph between key domains in Fig. 2c and d (*right*) shows that the I28A mutation dramatically weakens the interactions between the WW domain and PPIase core/α2-α3 helices, which indicates the fluctuation of the WW domain blocks the propagation of the allosteric signals from the WW to the PPIase core and α2-α3 helices. Although the WW domain is still partially connected to the α1-helix, the α1-helix cannot bridge to the catalytic loop, resulting in the breakdown of the pathway from the WW domain to the catalytic loop via the α1-helix (Fig. 2d, *right* and Fig. 3c).

### Allosteric effect of G93A amyotrophic lateral sclerosis-linked mutation on SOD1

Copper, zinc superoxide dismutase-1 (SOD1) is an oxidoreductase responsible for decomposing toxic superoxide radicals into molecular oxygen and hydrogen peroxide in two rapid steps by alternately reducing and oxidizing active-site copper^21^. The overall structure is composed of eight antiparallel β-strands, plus two loops forming an active site (Fig. 4a). The long active loop (residues 49-83) can be divided into a dimerization subloop (DL), disulfide subloop (DSL), and a zinc-binding subloop (ZL). The small active loop is an electrostatic loop (EL) (residues 122-142) near the metal site^22^. A study of SOD1-linked neurodegenerative disorder amyotrophic lateral sclerosis (ALS) shows that the G93A mutation enforces the EL to move away from ZL, decreasing the Zn (II) affinity of the protein^23^, which affects the pathogenic process of SOD1-linked ALS^24^. Since the G93A mutation occurs away from the metal site (Fig. 4b), this process is allosteric.

**Fig. 4.**
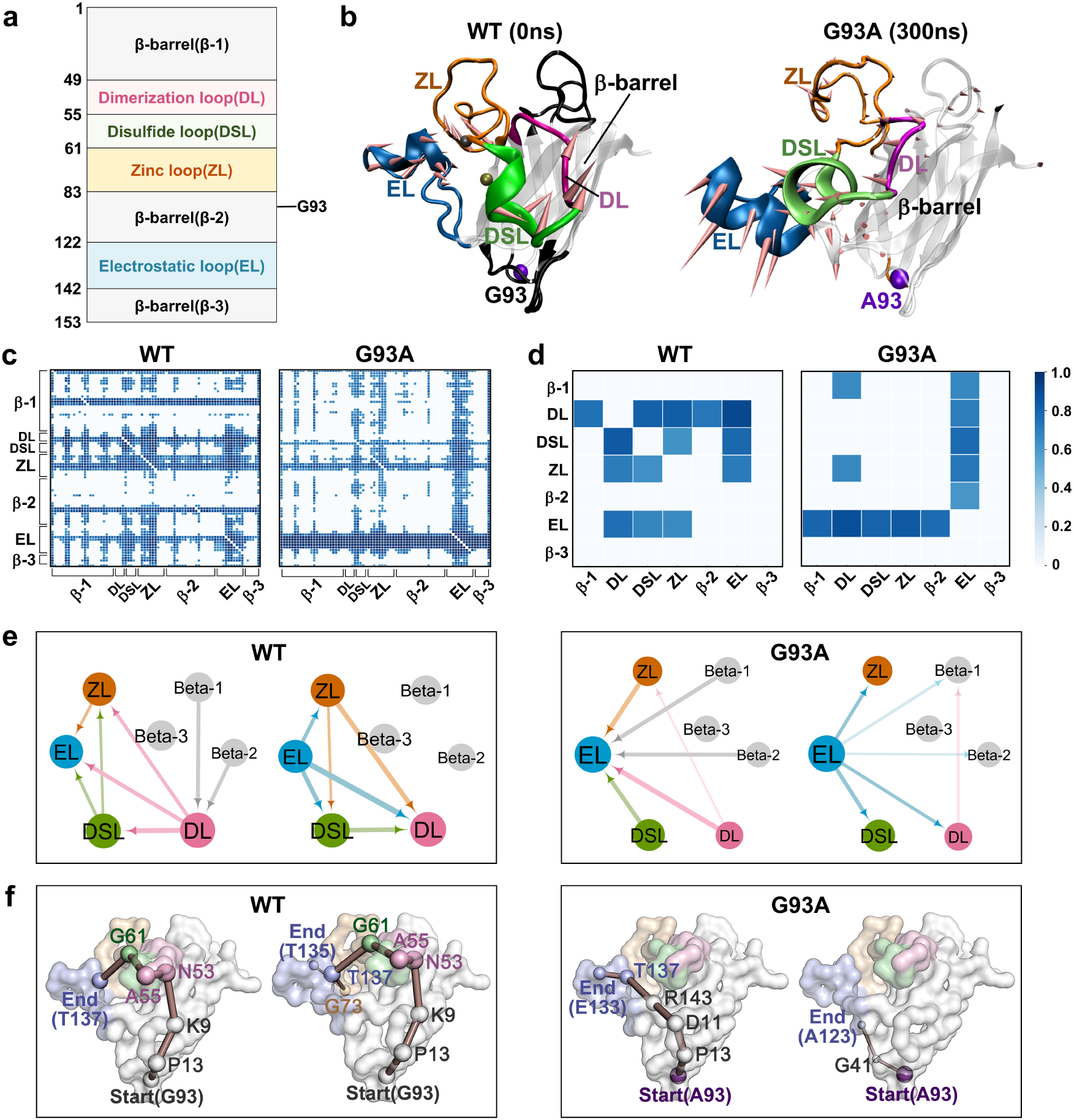
Change of domain interactions upon G93A mutation in SOD1. **a**, Domain partition of SOD1 protein including the position of G93A mutation. **b**, The initial structure of WT SOD1 and the G93A SOD1 structure at 300 ns, including a β-barrel (gray), dimerization subloop (DL) (pink), disulfide subloop (DSL) (green), zinc-binding subloop (ZL) (orange), and an electrostatic loop (EL) (blue). The directions shown in the cartoon denote the motion mode of the protein. **c**, Distribution of learned edges between residues in the MD simulations of the WT (*left*) and G93A (*right*) SOD1. **d**, Distribution of learned edges between domains in the MD simulations of the WT (*left*) and G93A (*right*) SOD1. **e**, The interaction graph mapped from the learned edges for the WT (*left*) and G93A (*right*) SOD1. The size of a node represents the number of edges that directly connect to the node. The thickness of an edge represents the strength of the interaction. The arrowhead denotes the directionality of a learned edge. **f**, The pathways from the G93 through residues in the β-barrel and residues in the long active loop to the EL loop for the WT SOD1 (*left*), and the pathways from the A93 through residues in the β-barrel to the EL loop for the G93A SOD1 (*right*). The size of a node represents the number of edges that directly connect to the node. The thickness of an edge represents the strength of the interaction. We used G93/A93 as the starting point and residues in the EL as the ending points to present the pathways.

We performed MD simulations for WT and G93A SOD1 to generate trajectories for learning the interactions in SOD1 (see Methods for detail). The RMSF values (Fig. S4) show that the WT and G93A SOD1 exhibit high flexibility in the long active loop and the EL. In particular, the EL of the G93A SOD1 becomes more flexible than that of the WT SOD1. Correspondingly, the motion mode reveals that the G93A mutation induces the EL far away from the metal site, while the EL of the WT SOD1 can be stabilized in the proximity of the metal site (Fig. 4b). Then, we ran the NRI model on the trajectories and compared the performance of motion reconstruction. From the ground truth and reconstructed trajectories (Supplementary Video 2), the RMSF values (Fig. S5), and the MSE values (Table S1), we find that the NRI model can reconstruct the dynamics many time steps based on the interaction graph learned by the encoder.

The interacting domains mapped from the learned graph show that the long active loop (DL, DSL, and ZL) and the small active loop (EL) can interact with each other closely in the WT SOD1 (Fig. 4c, d, and e, *left*). A close look at the learned edges graph in Fig. 4c, *left* reveals that the long and small active loops also connect to the residues in β-barrel, which are responsible for the close state of the EL. Moreover, the pathways in WT SOD1 elucidate that the communication can start from G93 through the long active loop to the EL (Fig. 4f, *left* and Table S3). In contrast, during the EL opening induced by the G93A mutation, the inner connections originally between the long active loop and the β-barrel in the WT SOD1 are almost broken, making the EL detached from the metal site (Fig. 4c, d, and e, *right*). Then the allosteric pathways emanate from the A93 no longer propagate through the long active loop, but directly through the residues in β-barrel to the EL (Fig. 4f, *right* and Table S3).

Comparing the interaction graph to the result of RMSF (Fig. 4 c, d, and Fig. S4) shows that the highly flexible domains usually play an essential role in a dynamic interacting system. For example, in the movement of the WT SOD1, more edges are connected to the long and small active loops instead of the β-barrel. Correspondingly, the RMSF values of the active loops are higher than those of the β-barrel. For the G93A SOD1, the whole protein structure becomes more flexible. In particular, since the EL’s open trend induces a significant increase in flexibility, the interaction of other domains is weakened. In addition, we find that the G93A mutation makes the A93(O)-L38(N) distance increase, resulting in a decrease in hydrogen bond interactions (Fig. S6a and Table S4). The increased distance in A93(O)-L38(N) weakened the hydrogen bond interactions between β-barrel and active loops (Figs. S6b-i, S7, and Table S4), making the SOD1 structure looser compared to the WT SOD1. Also, the overall dimensions of the proteins calculated by the radius of gyration (Rg) demonstrate the decreased protein compactness (Fig. S8) in the G93A SOD1, compared to that in the WT SOD1.

### Mechanism of oncogenic mutations activating MEK1

Mitogen-activated protein kinase kinase (MAPKK, also known as MEK) acts as an integration point in the RAS-RAF-MEK-ERK mitogen-activated protein kinase (MAPK) signaling cascade^25^. The activation of MEK requires its phosphorylation by upstream kinases named RAF kinases^26^. The human MEK1 protein consists of a small N-terminal lobe (N-lobe) and a large C-terminal lobe (C-lobe)^27^. As shown in Fig. 5a and b, the small N-lobe is dominated by five antiparallel β-strands (core kinases domain-1) and two conserved αA/αC helices, in which the αC-helix is critical in the regulation of MEK1 activity^28^. The active site of MEK1 is located at the interface of the N-lobe and C-lobe, binding to the substrate (such as ATP) or the competitive inhibitor atrial natriuretic peptide (ANP). The large C-lobe contains three core kinase domains, an activation segment, and a proline-rich loop, where the activation segment and proline-rich loop are crucial in regulating the activation of MEK1 and downstream ERKs in cells^28,29^. Recent studies reported that the E203K mutation can remotely affect the active site of MEK1 to increase the phosphorylation of ERK1/2^30^. Similarly, phosphorylation of Ser218 and Ser222 is also required for MEK1 activation to promote cell proliferation and transformation eventually leading to various human cancers^26,31^.

**Fig. 5.**
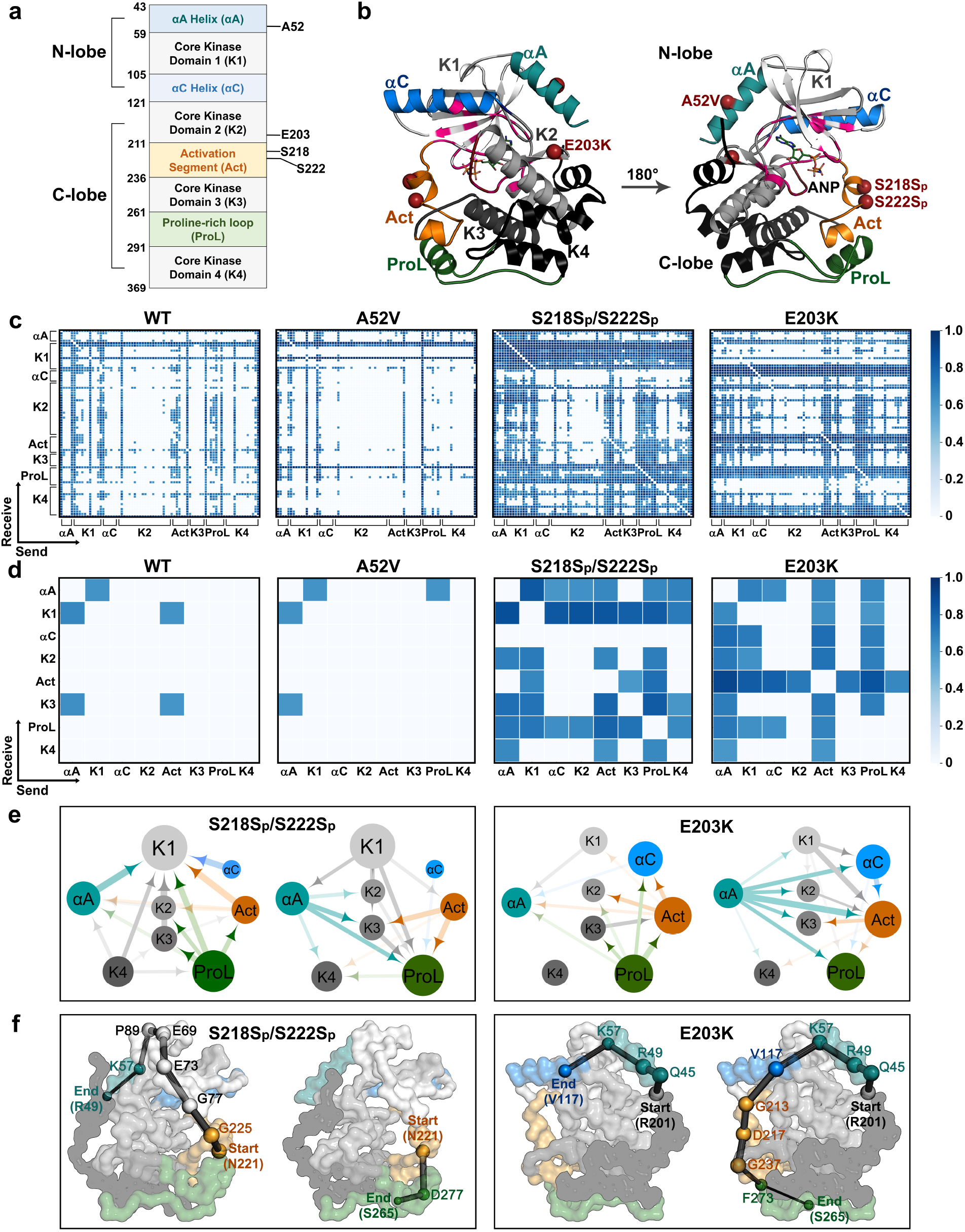
Change of domain communications upon active mutations in MEK1. **a**, Domain partition of MEK1 protein including the positions of mutations (A52V, S218Sp/S222Sp, and E203K). **b**, Two different views of MEK1 structure. The N-terminal lobe (N-lobe) contains one core kinase (gray) and two conserved α-helices (blue). The C-terminal lobe (C-lobe) contains three core kinase domains (gray and black), an activation segment (orange), and a proline-rich loop (green). **c**, Distribution of learned edges between residues in the MD simulations of WT, A52V, S218Sp/S222Sp, and E203K MEK1. **d**, Distribution of learned edges between domains in the MD simulations of WT, A52V, S218Sp/S222Sp, and E203K MEK1. **e**, The interaction graph mapped from the learned edges of active mutant MEK1. The size of a node represents the number of edges that directly connect to the node. The thickness of an edge represents the strength of the interaction. The arrowhead denotes the directionality of a learned edge. **f**, The allosteric pathways from N221 in the activation segment to the αA-helix and the proline-rich loop in the S218Sp/S222Sp MEK1 (*left*), and from R201 (near to E203K) to the αC-helix and the proline-rich loop in the E203K MEK1(*right*). The size of a node represents the number of edges that directly connect to the node. The thickness of an edge represents the strength of the interaction. We used N221 and R201 (near to E203K) as the starting points and residues in the αA, αC helices, and the proline-rich loop as the ending points to present the pathways.

To explore the allosteric effect of the mutation on MEK1, we performed MD simulations and analyses for two nonactive MEK1 (WT and A52V)^30^, two active forms (mutation E203K^30^, and a phosphorylated MEK1, where both Ser218 and Ser222 are phosphorylated)^26,31^. The secondary structure changes (Fig. S9) show that the activation segment experiences a helix-to-loop transition in the active MEK1 (E203K and phosphorylated Ser218/222). In contrast, this segment’s helix content in the WT and A52V MEK1 increased significantly compared to the active MEK1. The principal component analysis (Fig. S10) reflects the activation segment’s open trend in the active MEK1.

The above analysis only shows the changes in the dynamic motions of MEK1. To explore the interaction patterns in the MEK1 motions, the NRI model was applied to learn the trajectories. The motion reconstruction results show that the trajectory reconstructed by the NRI model almost coincides with the ground-truth trajectory (Fig. S11, Supplementary Video 3, and Table S1), indicating the high degree of confidence of the learned interaction graph. As shown in the learned interaction graph of nonactive MEK1 (WT and A52V) (Fig. 5c and d), few interactions occur between the domains. In contrast, the αA-helix, core kinase domain-1, activation segment, and the proline-rich loop of phosphorylated MEK1 can interact with other domains, which indicates that these domains drive the slow motion in the activation of phosphorylated MEK1 (Fig. 5c and d).

We mapped the graph of the phosphorylated MEK1 as the interacted domains (Fig. 5e, *left*) and calculated the allosteric pathways (Fig. 5f, *left* and Table S5). Interestingly, four domains (αA-helix, core kinase domain-1, activation segment, and proline-rich loop) form an interaction pattern, in which the activation segment connects all the way to the core kinase domain-1 and αA-helix, which may affect the binding affinity of ANP in the active pocket. Meanwhile, the activation segment can also connect to the proline-rich loop, which may activate downstream ERKs in cells. Then, we applied the NRI model to learn the inner-domain correlation from the dynamic motion of E203K MEK1. A closer look at the learned graph reveals that similar to the phosphorylated MEK1, the active mutation (E203K) strengthens the interactions between the activation segment/proline-rich loop and the rest of MEK1 (Fig. 5c, d, and e). From the allosteric pathways starting with R201 (Fig. 5f, *right* and Table S5), we find that the activation segment has a great effect on transiting message from R201 (near to E203K) to the proline-rich loop. For the effect of the E203K mutation on the αC-helix, the communication propagates through the αA-helix to the αC-helix. Hence, phosphorylated Ser218/222 and E203K mutations have a similar spatial effect on the MEK1 structure, i.e., the activation segment as a “messenger” can interact with the N-lobe and proline-rich loop in the dynamic and both can affect the N-lobe (active site) and the communication in the proline-rich loop to regulate the binding to the downstream substrates.

## Discussion

This study applied a GNN-based NRI model to analyze latent interactions between residues from reconstructing MD trajectories of proteins. We carried out three case studies to explore the allosteric long-range interactions for the Pin1, SOD1, and MEK1 systems. We have demonstrated that the NRI model can effectively generate the interaction graphs related to protein’s slow-motion through an embedding of reconstructing MD trajectories, and the shortest pathways between the allosteric site and the active site in the interaction graphs can reveal the pathways mediating allosteric communications. Although many methods have applied graph theory to model allosteric communication in protein, they are mostly based on the static crystal structures of proteins only. These methods are ineffective in modeling allostery, which is dynamic in nature. One major advance in our modeling allostery is the use of information from MD simulations. Notably, we take a major step in recognizing allostery related long-range interactions by adding an instantaneous velocity feature to each node over the continuous snapshots, which can explain the allosteric communication related to biomolecular slow-motion. The allosteric pathways derived from the shortest paths provide valuable information when considering protein design. It may be possible to mutate the residues in the allosteric pathways to alter the biological functions and regulatory properties of proteins. Future research directions may include a longer MD simulation or experimental works to validate the predictions from the NRI model.

With regard to traditional analysis methods applied to MD trajectories, PCA obtains principal components representing the movement patterns determined in MD simulations, and the cross-correlation analysis is a measurement that tracks the movements over time of two variables to determine the degree of linear correlation between them. However, these two classical methods are limited to linear correlations mainly associated with harmonic/isotropic local torsional motion that occur on relatively short-time scales. It is well known that dynamic biomolecules often undergo large-scale structural changes during their biological functions. Due to large energy barriers, the conformational changes of biomolecules usually occur in the millisecond or longer time scales in nonlinear fashions. The NRI model is a powerful tool to model nonlinear relationships by introducing a nonlinear activation function. Not only for the allosteric regulation, many other biological and pharmaceutical processes, such as protein folding/unfolding, protein activation, or drug molecule binding targets can also be formulated as a dynamic interaction graph by the NRI model. In particular, the NRI model is appealing when probing the non-periodic biomolecular motion. Unlike the periodic physical movement where the interactions do not change over time, proteins during performing functions are often accompanied by considerable conformation and interaction changes. We believe the NRI model can be developed into a powerful tool for these problems in general.

## Methods

### NRI model

The original NRI model^15^ consists of two co-training parts: an encoder to predict the interaction given the dynamic system’s trajectories, and a decoder to predict the trajectories of the dynamic system given the interaction graph. The NRI model simultaneously learns the edge values and reconstructs the trajectories of the dynamic system in an unsupervised manner based on an unknown graph *z* (where *z*_*ij*_ represents the discrete attribute value of the edge between nodes *v*_*i*_ and *v*_*j*_). The input consists of *N* nodes. The feature vector (position and velocity in the dimensions of x, y, and z) of node *v*_*i*_ (6-dimention for each node) is denoted as 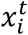 at time *t*. All *N* nodes’ feature set is denoted as 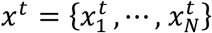. The trajectory of node *i* is denoted as 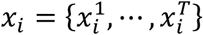, where *T* is the number of time steps. Finally, all trajectory data are recorded as *x* = {*x*^1^, ⋯ , *x*^*T*^}. The structure of the model is presented in Fig. 1.

As shown in the left of Fig. 1, the encoder

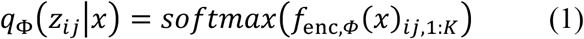

infers the discrete categorical variable *z*_*ij*_ based on the input trajectories *x*_1_, ⋯ , *x*_*K*_, in which *f*_*enc,ϕ*_(*x*) is a GNN performed on the fully connected networks (without self-connection) to predict the latent graph structure. The encoder runs two rounds of node-to-edge (*ν* → *e*) and an edge-to-node (*e* → *ν*) message passing. The node-to-edge operation generates the edge features connecting the node embeddings and the edge-to-node operation can aggregate the message of edge embeddings from all incoming edges. Since the graph is fully connected, each node obtains a message from the entire graph. Finally, all messages pass from nodes to edges. In our implementation model, every message passing operation is performed by a 2-layer multilayer perceptron (MLP)^15^.

The distribution of *z*, *q*_*ϕ*_(*z*|*x*), is learned from the encoder. Then the sampling is performed to generate *z*_*ij*_ only available in *K* edge type. We sampled from a continuous approximation of the discrete distribution and used the repramatrization to obtain gradients from this approximation^32^, i.e.

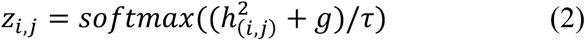

where *g* ∈ ℝ^*K*^ is an independent and uniformly distributed vector from the Gumbel distribution (0,1), *τ* (softmax temperature) represents the smoothness of sampling, and the distribution tends to become one-hot samples when *τ* → 0.

The decoder

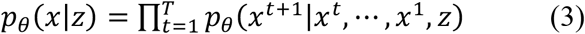

reconstructs the dynamic systems *p*_*θ*_(*x*^*t*+1^|*x*^*t*^, ⋯ , *x*^1^, *z*) with a GNN given the latent graph structure *z*. A recurrent decoder with a GRU unit^33^ is required to model *p*_*θ*_(*x*^*t*+1^|*x*^*t*^, ⋯ , *x*^1^, *z*). The decoder runs multiple GNNs in parallel to the encoder. In the node-to-edge (*v* → *e*) message passing, the input is the recurrent hidden state at the previous time step. The hidden state of an edge is determined by the hidden state of its connecting nodes and it allows the message at each time step passing through the hidden state. Thus, the prediction at *t* + 1 is based not only on the previous time step but also on messages from all previous time steps. In the edge-to-node (*e* → *v*) message passing, the concatenation of the aggregated messages, the current input, and the previously hidden state, is denoted as the input of GRU update to generate the hidden state at the next time step. Then the value observed previously and the hidden state at the current time step are used to predict the state’s distribution (position and velocity) in future time steps.

This model is formalized as a variational autoencoder (VAE)^34,35^ that maximizes the Evidence Lower Bound (ELBO):

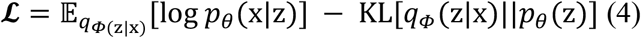

The ELBO objective has two terms, namely the reconstruction error 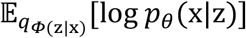 and KL divergence KL[*q*_*ϕ*_(z|x)∥*p*_*θ*_(z)], in which the encoder *q*_*ϕ*_(z|x) returns a factorized distribution of *zij,* and one-hot representation of the *K* interaction types (*K*=4 presents edges/no edge) is used on *z*_*ij*_. *p*_*θ*_(*x*|*z*) represents the decoder that reconstructs the dynamic systems given the distribution of *z*_*ij*_. The autoencoder maps the input *X* to a feature space, and then map from this feature space back to the input space to minimize the reconstruction error. The reconstruction error is calculated through:

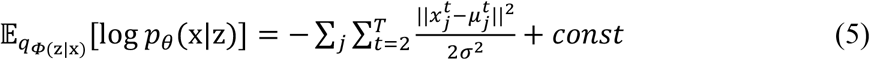

and the KL divergence is the sum of entropies and a constant:

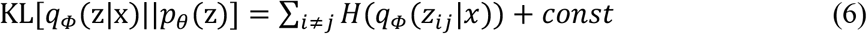

The whole training process was carried out as follows: (*i*) We first performed the encoder to calculate *q*_ϕ_(*z*_*ij*_,*x*) given a training MD trajectory *X* , (*ii*) we then sampled *z*_*ij*_ from a continuous approximation of the discrete distribution, and (*iii*) we finally ran the decoder to reconstruct the interacting dynamics *p*_*θ*_(*x*^*t*+1^|*x*^*t*^, ⋯ , *x*^1^, *z*) for Pin1, SOD1 and MEK1 systems.

### Simulation data

#### 1. Preparation of protein structures

The crystal structures for the three systems (Pin1, SOD1, and MEK1) were retrieved from the Protein Data Bank (www.rcsb.org). The apo Pin1 structure was obtained from PDB 3TDB^17^. To obtain the Pin1-FFpSPR complex, we docked the substrate (FFpSPR) into the WW-domain of apo Pin1 using Autodock 4.2^36^. Missing residues of the protein (residues 39-50) in the inter-domain linker were modeled by SWISS-MODEL^37^. The SOD1 and MEK1 structures were taken directly from PDB 2C9V^22^ and 3SLS^27^. The structures of corresponding mutations were also constructed using the SWISS-MODEL.

#### 2. Conventional molecular dynamics (cMD) simulations

The cMD simulations of four MEK1 structures (WT, A52V, E203K, and phosphorylated MEK1) were performed using GROMACS 5.1.4 package^38^ with Gromos 53A6 force field^39^. All complexes of MEK1 structures and an inhibitor (ANP) were performed in a periodic boundary box with the SPC water model^40^. And chloride or sodium ions were randomly added to this box to neutralize the systems. In addition, energy minimization was performed using the steepest descent method to obtain the energy-minimized initial structure for the next simulations. Subsequently, 100 ps of NVT (Berendsen temperature coupled with constant particle number, volume, and temperature)^41^ and 100 ps of NPT (Parrinello-Rahman pressure coupled with constant particle number, pressure, and temperature)^41^ were performed to maintain the stability of the system (300 K, 1 bar). The coupling constants for temperature and pressure were set at 0.1 and 2.0 ps, respectively. Long-range electrostatic interactions were described using the particle mesh Ewald algorithm with an interpolation order of 4 and a grid spacing of 1.6 Å^42^. Van der Waals interactions were calculated according to the cutoff value of 12 Å. All bond lengths were constrained using the LINear Constraint Solver (LINCS) algorithm^42^. After stabilizing all thermodynamic properties, the molecular systems were simulated for 200 ns with a time interval of 2 fs, whereas the coordinates for all models were stored every 2 ps.

#### 3. Gaussian accelerated molecular dynamics (GaMD) simulation

GaMD simulation is an enhanced sampling technique by adding a harmonic boost potential to smoothen the system’s potential energy surface^43,44^. To enhance the conformational sampling of Pin1 and SOD1 structures, GaMD simulations were performed on the apo Pin1, FFpSPR bound Pin1, FFpSPR bound I28A Pin1, WT SOD1, and G93A SOD1 structures. For each of the systems, the graphics processing unit (GPU) version of AMBER18 was applied to perform GaMD simulation^45^. The GaMD simulation has five stages: (*i*) conventional MD preparatory stage for the equilibration of the system, (*ii*) conventional MD stage to collect potential statistics for calculating the GaMD acceleration parameters, (*iii*) GaMD pre-equilibration stage with boost potential, (*iv*) GaMD equilibration stage to update the boost parameters, (*v*) multiple independent GaMD production runs with randomized initial atomic velocities^43^. The molecular systems were simulated 200 ns for Pin1 system and 300 ns for SOD1 system, with a time interval of 2 fs. The GaMD simulation trajectories were analyzed using CPPTRAJ^46^ and VMD^47^ for RMSF calculation, secondary structure analysis, principal component analysis, and hydrogen bond calculation.

### NRI model construction details

All NRI trainings were performed using Adam optimizer^48^ with a learning rate of 0.0005 and a batch size of 1, decayed by a factor of 0.5 every 200 epochs. The concrete distribution was used with *τ* = 0. During testing, we replaced the concrete distribution with a categorical distribution to obtain discrete latent edge types. All experiments were run for 500 training epochs. The discrete samples were used in the training forward pass. We saved model checkpoints after every epoch whenever the validation set performance improved and loaded the best performing model for the test set evaluation. We used a standard Nvidia GeForce GTX 1080Ti GPU card and a Core solo CPU to train our models. Each CPU was allocated 48G memory. The training time for one experiment took about 5 hours.

We used the MD trajectories to generate the input data for the next training, validation, and test. The dataset of the Pin1 system has a total size of 2000 frames for 73 Cα atoms each. The dataset of the SOD1 system has 3000 frames for 77 Cα atoms each. The dataset of the MEK1 system has 2500 frames for 76 Cα atoms each. We normalized the position and velocity features to the maximum absolute value of 1. The overall input/output dimension of the model is 6 (3D position and velocity). Training, validation, and test samples each contain 50 frames uniformly extracted from each trajectory. For the training part, the model receives a ground truth input in each timestep. The dynamics for our three systems change considerably over time, the protein conformations in the early stage of the simulation are quite different from that in the end stage. Therefore, in the test of the experiment, we fed in the 50 timesteps as ground truth to the encoder and then reconstructed these timesteps. All experiments used MLP encoder and recurrent neural network (RNN) decoder to have a comparable capacity to the full graph model. The first edge type is “hard-coded” as non-edge (no messages are passed along this type). The basic building block of the MLP encoder is a 2-layer MLP with a hidden and output dimension of 256, with batch normalization, dropout, and ELU activations. The RNN decoder adds a GRU style update to the single-step prediction. Given the interaction graph learned from the NRI model, we took the allosteric site as the starting point and the active site as the terminal point to calculate the shortest pathways using Dijkstra’s algorithm^49^.

## Acknowledgement

This work was supported by the China Scholarship Council to JZ and the US National Institutes of Health [R35-GM126985] to DX.

## References

Altis, A., Nguyen, P. H., Hegger, R. & Stock, G. Dihedral angle principal component analysis of molecular dynamics simulations. The Journal of chemical physics 126, 244111, doi:10.1063/1.2746330 (2007).

Hünenberger, P. H., Mark, A. E. & van Gunsteren, W. F. Fluctuation and cross-correlation analysis of protein motions observed in nanosecond molecular dynamics simulations. Journal of molecular biology 252, 492–503, doi:10.1006/jmbi.1995.0514 (1995).

Li, D. W., Meng, D. & Brüschweiler, R. Short-range coherence of internal protein dynamics revealed by high-precision in silico study. Journal of the American Chemical Society 131, 14610–14611, doi:10.1021/ja905340s (2009).

Lange, O. F. & Grubmüller, H. Generalized correlation for biomolecular dynamics. Proteins 62, 1053–1061, doi:10.1002/prot.20784 (2006).

Long, S. & Tian, P. Nonlinear backbone torsional pair correlations in proteins. Scientific reports 6, 34481, doi:10.1038/srep34481 (2016).

Atilgan, A. R., Akan, P. & Baysal, C. Small-world communication of residues and significance for protein dynamics. Biophysical journal 86, 85–91, doi:10.1016/s0006-3495(04)74086-2 (2004).

Sethi, A., Eargle, J., Black, A. A. & Luthey-Schulten, Z. Dynamical networks in tRNA:protein complexes. Proc. Natl. Acad. Sci. U. S. A. 106, 6620–6625, doi:10.1073/pnas.0810961106 (2009).

Wu, Z. et al. A Comprehensive Survey on Graph Neural Networks. IEEE Transactions on Neural Networks and Learning Systems, 1–21, doi:10.1109/TNNLS.2020.2978386 (2020).

A, J. W. et al. Inductive Inference of Gene Regulatory Network Using Supervised and Semi-supervised Graph Neural Networks. (2020).

Wang, J. et al. scGNN: a novel graph neural network framework for single-cell RNA-Seq analyses. bioRxiv, 2020.2008.2002.233569, doi:10.1101/2020.08.02.233569 (2020).

Hoshen, Y. VAIN: Attentional Multi-agent Predictive Modeling. (2017).

Guttenberg, N., Virgo, N., Witkowski, O., Aoki, H. & Kanai, R. Permutation-equivariant neural networks applied to dynamics prediction. (2016).

Mnih, V., Heess, N., Graves, A. & Kavukcuoglu, K. J. a. e.-p. Recurrent Models of Visual Attention. arXiv:1406.6247 (2014). <https://ui.adsabs.harvard.edu/abs/2014arXiv1406.6247M>.

van Steenkiste, S., Chang, M., Greff, K. & Schmidhuber, J. J. a. e.-p. Relational Neural Expectation Maximization: Unsupervised Discovery of Objects and their Interactions. arXiv:1802.10353 (2018). <https://ui.adsabs.harvard.edu/abs/2018arXiv180210353V>.

Kipf, T., Fetaya, E., Wang, K.-C., Welling, M. & Zemel, R. Neural Relational Inference for Interacting Systems. (2018).

Gunasekaran, K., Ma, B. & Nussinov, R. Is allostery an intrinsic property of all dynamic proteins? Proteins 57, 433–443, doi:10.1002/prot.20232 (2004).

Zhang, M. et al. Structural and Kinetic Analysis of Prolyl-isomerization/Phosphorylation Cross-Talk in the CTD Code. ACS Chemical Biology 7, 1462–1470, doi:10.1021/cb3000887 (2012).

Yaffe et al. Sequence-specific and phosphorylation-dependent proline isomerization: A potential mitotic regulatory mechanism. (1997).

Namanja, A. T. et al. Stereospecific gating of functional motions in Pin1. 108, 12289–12294 (2011).

Wilson, K. A., Bouchard, J. J. & Peng, J. W. Interdomain interactions support interdomain communication in human Pin1. Biochemistry 52, 6968–6981, doi:10.1021/bi401057x (2013).

Hart et al. A Structure-Based Mechanism for Copper-Zinc Superoxide Dismutase. (1999).

Strange, R. W. et al. Variable metallation of human superoxide dismutase: atomic resolution crystal structures of Cu-Zn, Zn-Zn and as-isolated wild-type enzymes. 356, 1152–1162 (2006).

Kayatekin, C., Zitzewitz, J. A. & Matthews, C. R. Zinc binding modulates the entire folding free energy surface of human Cu,Zn superoxide dismutase. Journal of molecular biology 384, 540–555, doi:10.1016/j.jmb.2008.09.045 (2008).

Smith, A. P. & Lee, N. M. Role of zinc in ALS. Amyotrophic lateral sclerosis : official publication of the World Federation of Neurology Research Group on Motor Neuron Diseases 8, 131–143, doi:10.1080/17482960701249241 (2007).

Kolch, W. Coordinating ERK/MAPK signalling through scaffolds and inhibitors. Nature reviews. Molecular cell biology 6, 827–837, doi:10.1038/nrm1743 (2005).

Shi, H., Kong, X., Ribas, A. & Lo, R. S. Combinatorial treatments that overcome PDGFRβ-driven resistance of melanoma cells to V600EB-RAF inhibition. Cancer research 71, 5067–5074, doi:10.1158/0008-5472.Can-11-0140 (2011).

Meier, C. et al. Engineering human MEK-1 for structural studies: A case study of combinatorial domain hunting. 177, 329–334 (2012).

Hanks, S. K. & Hunter, T. The eukaryotic protein kinase superfamily: kinase (catalytic) domain structure and classification1. 9, 576–596, doi:https://doi.org/10.1096/fasebj.9.8.7768349 (1995).

Dang, A., Frost, J. A. & Cobb, M. H. The MEK1 proline-rich insert is required for efficient activation of the mitogen-activated protein kinases ERK1 and ERK2 in mammalian cells. The Journal of biological chemistry 273, 19909–19913, doi:10.1074/jbc.273.31.19909 (1998).

Gao, J. et al. 3D clusters of somatic mutations in cancer reveal numerous rare mutations as functional targets. 9, 4 (2017).

Atefi, M. et al. Reversing melanoma cross-resistance to BRAF and MEK inhibitors by co-targeting the AKT/mTOR pathway. 6, e28973 (2011).

Jang, E., Gu, S. & Poole, B. J. a. e.-p. Categorical Reparameterization with Gumbel-Softmax. arXiv:1611.01144 (2016). <https://ui.adsabs.harvard.edu/abs/2016arXiv161101144J>.

Cho, K. et al. Learning phrase representations using RNN encoder-decoder for statistical machine translation. (2014).

Kingma, D. P. & Welling, M. J. a. e.-p. Auto-Encoding Variational Bayes. arXiv:1312.6114 (2013). <https://ui.adsabs.harvard.edu/abs/2013arXiv1312.6114K>.

Jimenez Rezende, D., Mohamed, S. & Wierstra, D. J. a. e.-p. Stochastic Backpropagation and Approximate Inference in Deep Generative Models. arXiv:1401.4082 (2014). <https://ui.adsabs.harvard.edu/abs/2014arXiv1401.4082J>.

Morris, G. M. et al. AutoDock4 and AutoDockTools4: Automated docking with selective receptor flexibility. Journal of computational chemistry 30, 2785–2791, doi:10.1002/jcc.21256 (2009).

Biasini, M. et al. SWISS-MODEL: modelling protein tertiary and quaternary structure using evolutionary information. 42, W252–W258 (2014).

Apol, E. et al. GROMACS USER MANUAL (Version 5.0-rc1). (2014).

Oostenbrink, C., Soares, T. A., van der Vegt, N. F. & van Gunsteren, W. F. Validation of the 53A6 GROMOS force field. European biophysics journal : EBJ 34, 273–284, doi:10.1007/s00249-004-0448-6 (2005).

Mark, P. & Nilsson, L. Structure and Dynamics of the TIP3P, SPC, and SPC/E Water Models at 298 K. The Journal of Physical Chemistry A 105, 9954–9960, doi:10.1021/jp003020w (2001).

Berendsen, H. J. C., Postma, J. P. M., van Gunsteren, W. F., DiNola, A. & Haak, J. R. Molecular dynamics with coupling to an external bath. The Journal of chemical physics 81, 3684–3690, doi:10.1063/1.448118 (1984).

Darden, T., York, D. & Pedersen, L. Particle mesh Ewald: An N⋅log(N) method for Ewald sums in large systems. The Journal of chemical physics 98, 10089–10092, doi:10.1063/1.464397 (1993).

Miao, Y. & McCammon, J. A. in Annual reports in computational chemistry Vol. 13 231–278 (Elsevier, 2017).

Miao, Y., Feher, V. A. & McCammon, J. A. Gaussian Accelerated Molecular Dynamics: Unconstrained Enhanced Sampling and Free Energy Calculation. Journal of chemical theory and computation 11, 3584–3595, doi:10.1021/acs.jctc.5b00436 (2015).

Lee, T.-S. et al. GPU-accelerated molecular dynamics and free energy methods in Amber18: performance enhancements and new features. 58, 2043–2050 (2018).

Roe, D. R. & Cheatham, T. E., 3rd. PTRAJ and CPPTRAJ: Software for Processing and Analysis of Molecular Dynamics Trajectory Data. Journal of chemical theory and computation 9, 3084–3095, doi:10.1021/ct400341p (2013).

Humphrey, W., Dalke, A. & Schulten, K. VMD: visual molecular dynamics. Journal of molecular graphics 14, 33-38, 27–38, doi:10.1016/0263-7855(96)00018-5 (1996).

Kingma, D. P. & Ba, J. J. a. e.-p. Adam: A Method for Stochastic Optimization. arXiv:1412.6980 (2014). <https://ui.adsabs.harvard.edu/abs/2014arXiv1412.6980K>.

Dijkstra, E. W. A note on two problems in connexion with graphs. Numerische Mathematik 1, 269–271, doi:10.1007/BF01386390 (1959).

