## supplemental figures and tables for "Neural relational inference to learn allosteric long-range interactions in proteins from molecular dynamics simulations"

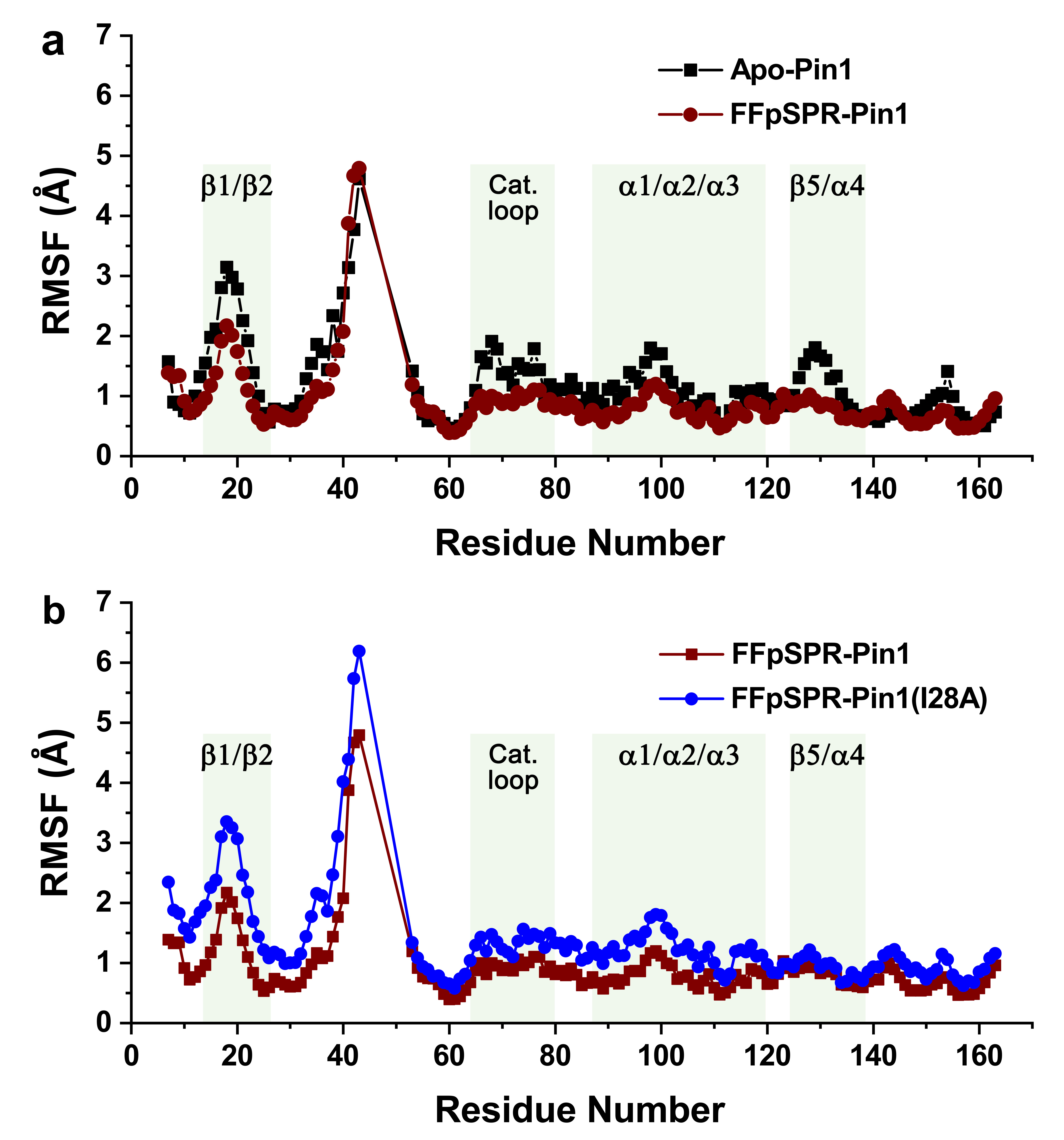

**Figure S1** | **a**, Comparison of RMSF values between apo Pin1 and FFpSPR bound Pin1. **b**, Comparison of RMSF values between FFpSPR bound Pin1 and FFpSPR bound Pin1 (I28A).

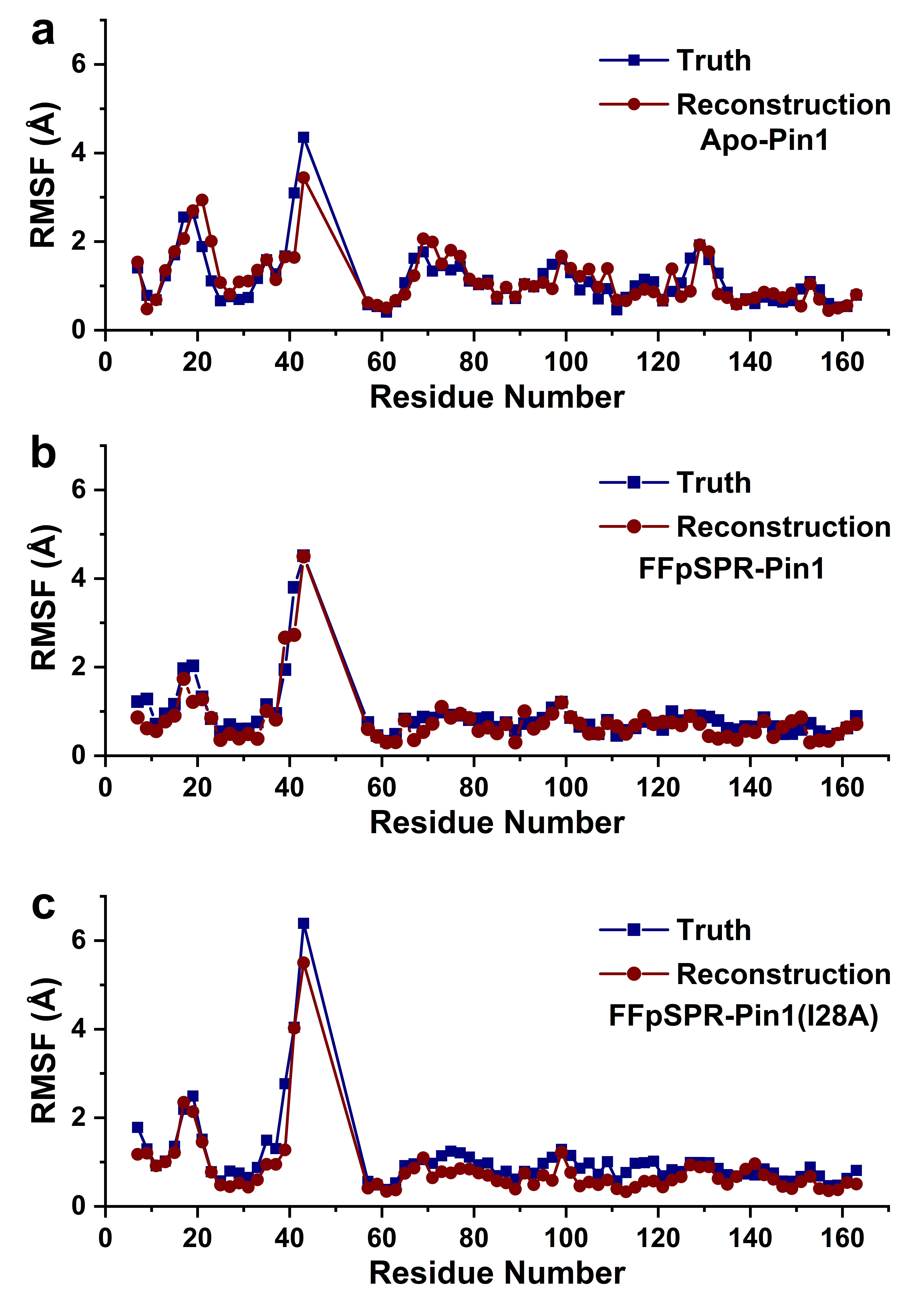

**Figure S2 |** Comparison of RMSF values between truth and reconstruction of trajectories for **a**, apo Pin1, **b**, FFpSPR bound Pin1, and **c**, FFpSPR bound Pin1 (I28A).

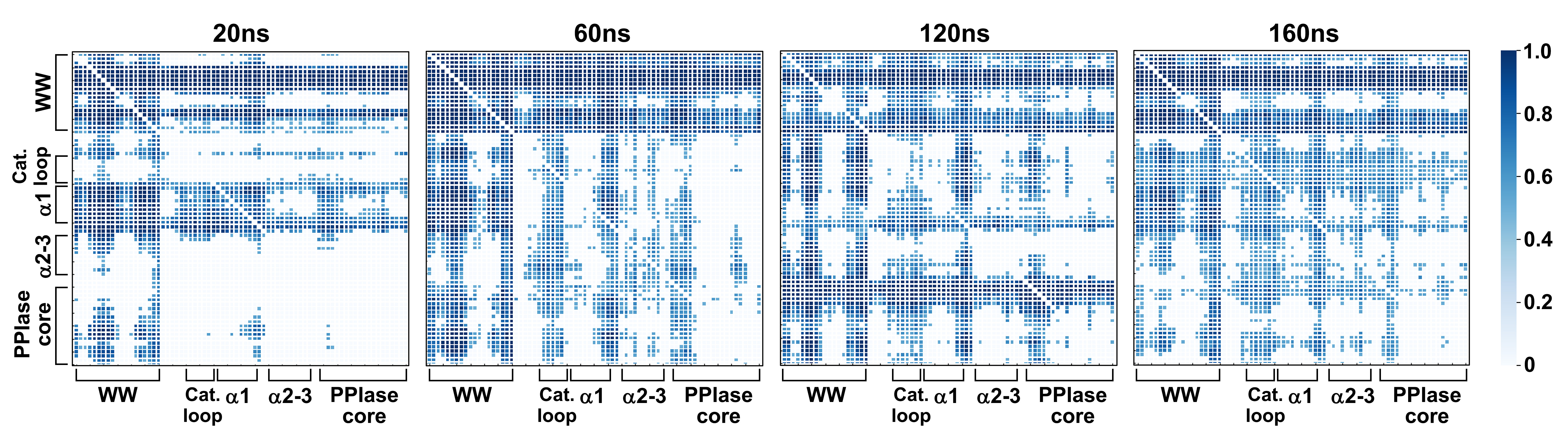

**Figure S3 |** Distribution of learned edges between amino acids in the MD simulations of different timesteps for FFpSPR bound Pin1.

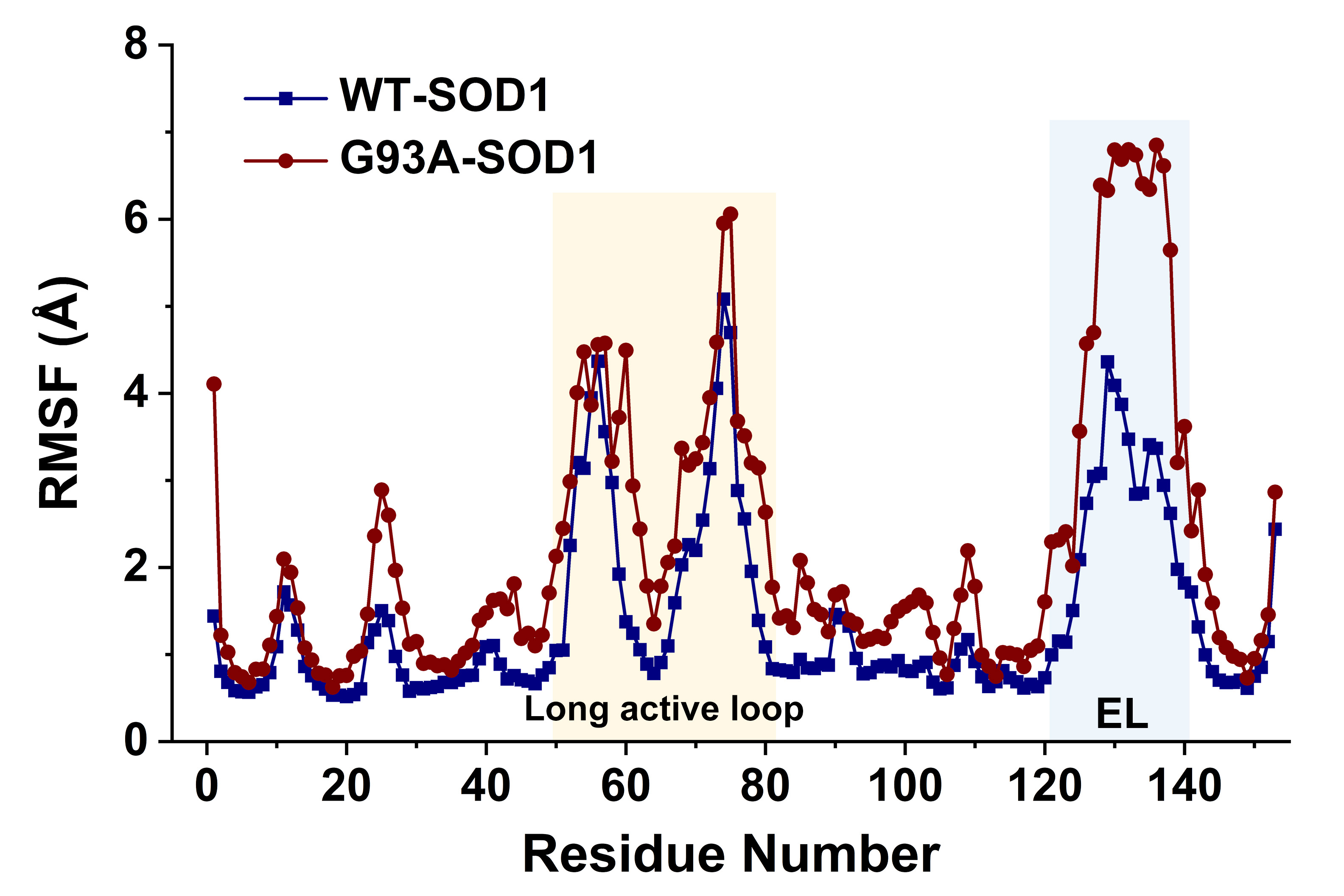

**Figure S4 |** RMSF plot of **a**, WT-SOD1 and **b**, G93A-SOD1 systems.

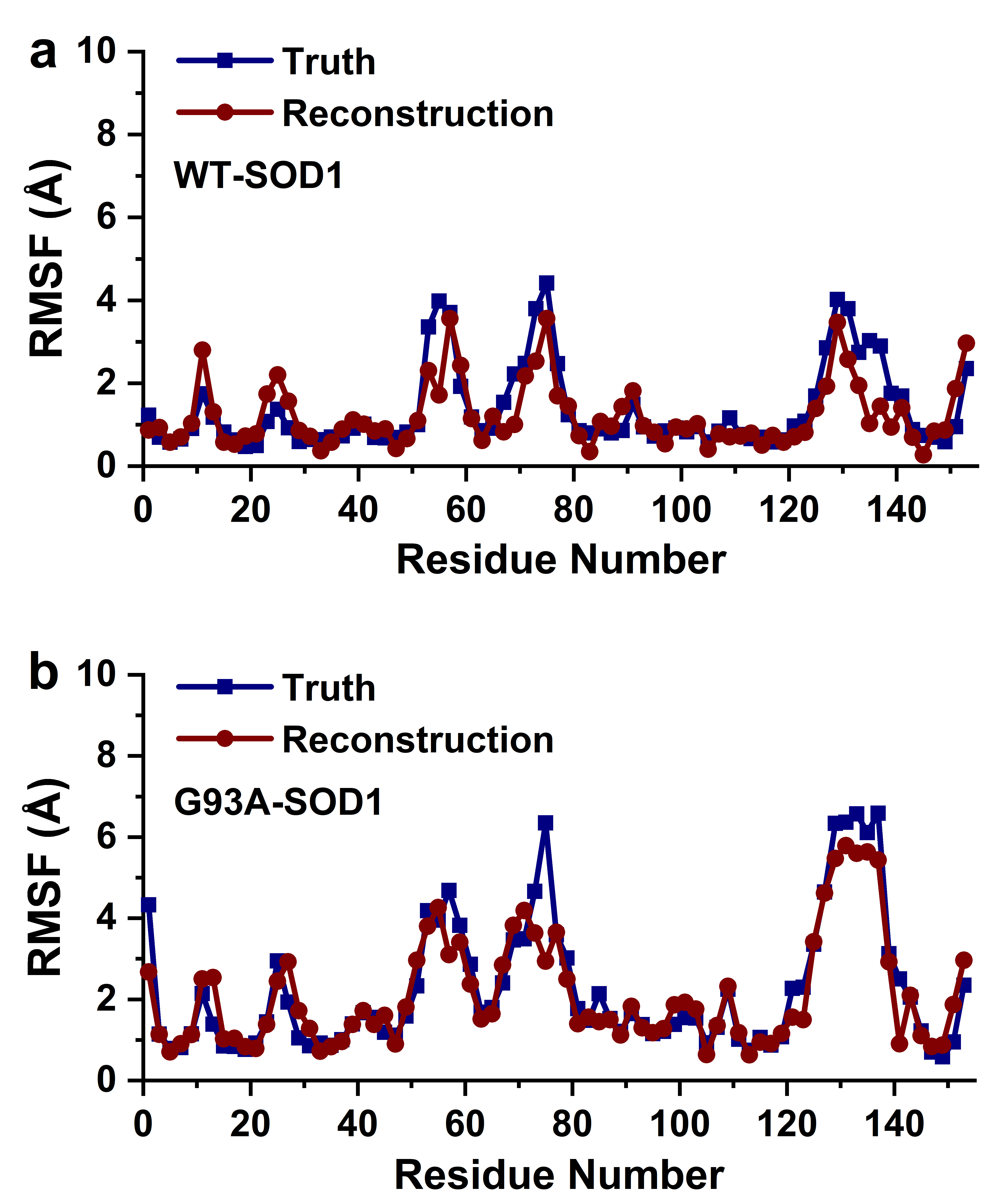

**Figure S5 |** Comparison of RMSF values between truth and reconstruction of trajectories for **a**, WT-SOD1 and **b**, G93A-SOD1.

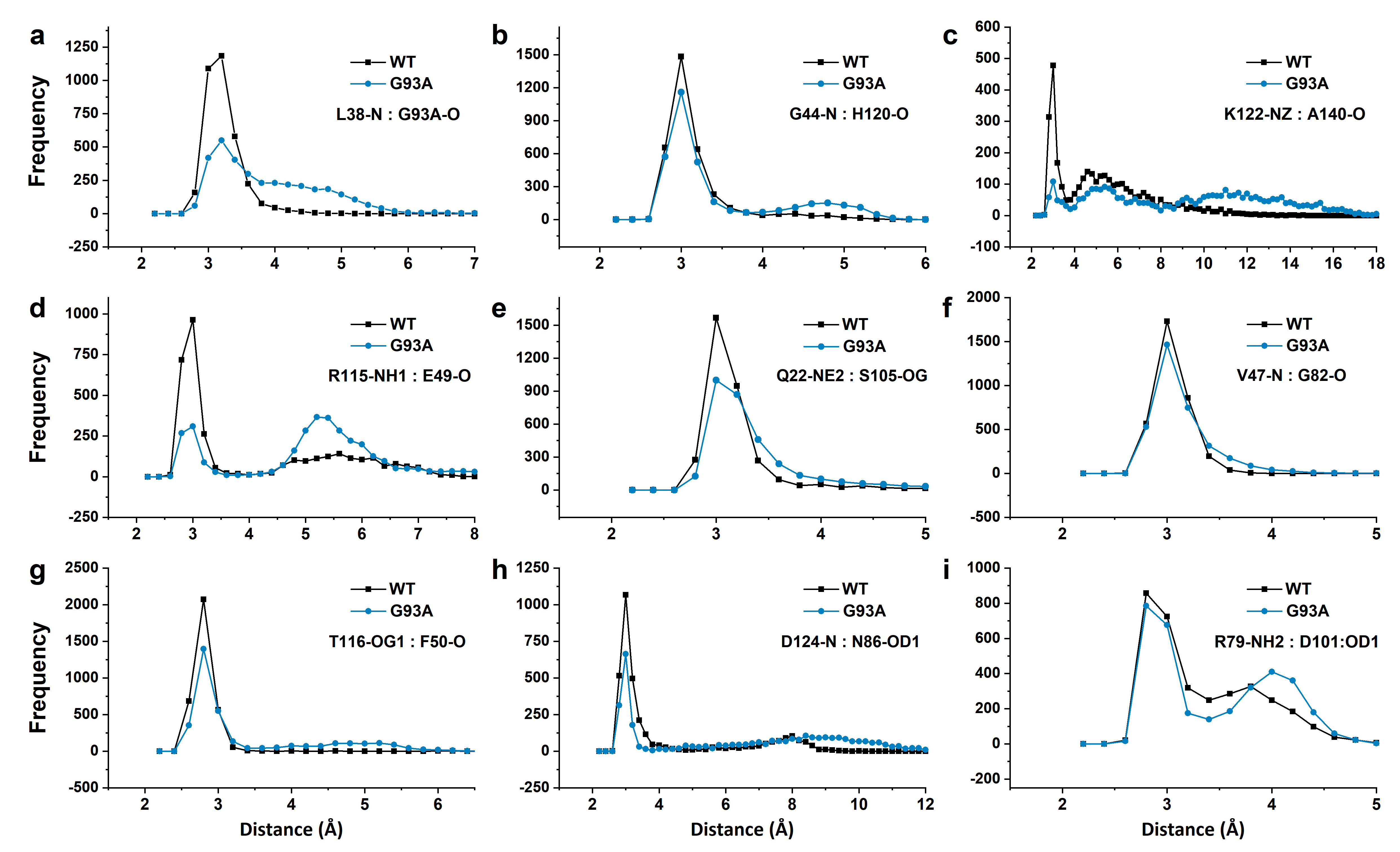

**Figure S6 |** Relevant distance distribution between atoms that form hydrogen bond interactions in the trajectories of WT-SOD1 (black) and G93A-SOD1 (blue).

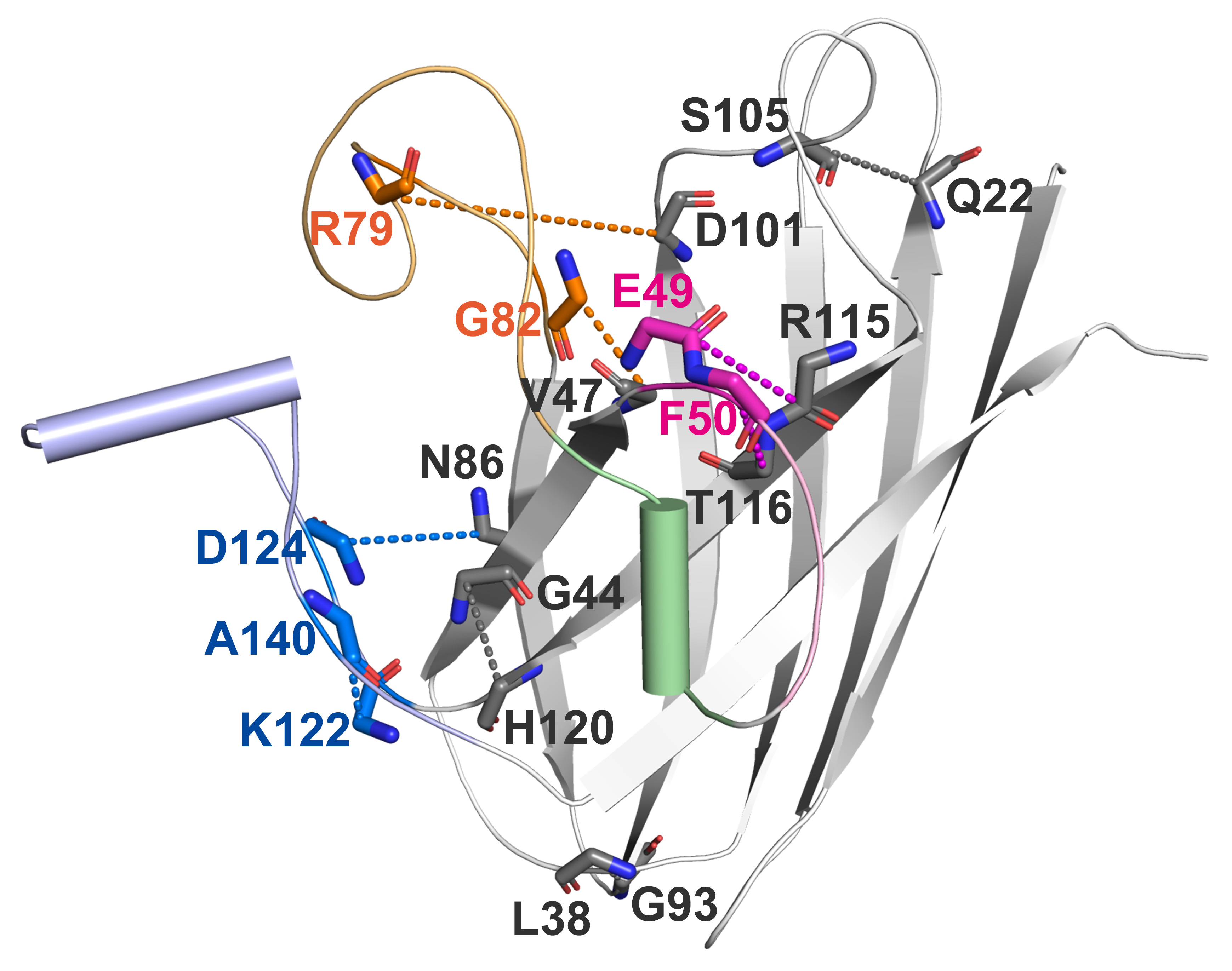

**Figure S7 |** SOD1 structure containing residues that form hydrogen bond interactions.

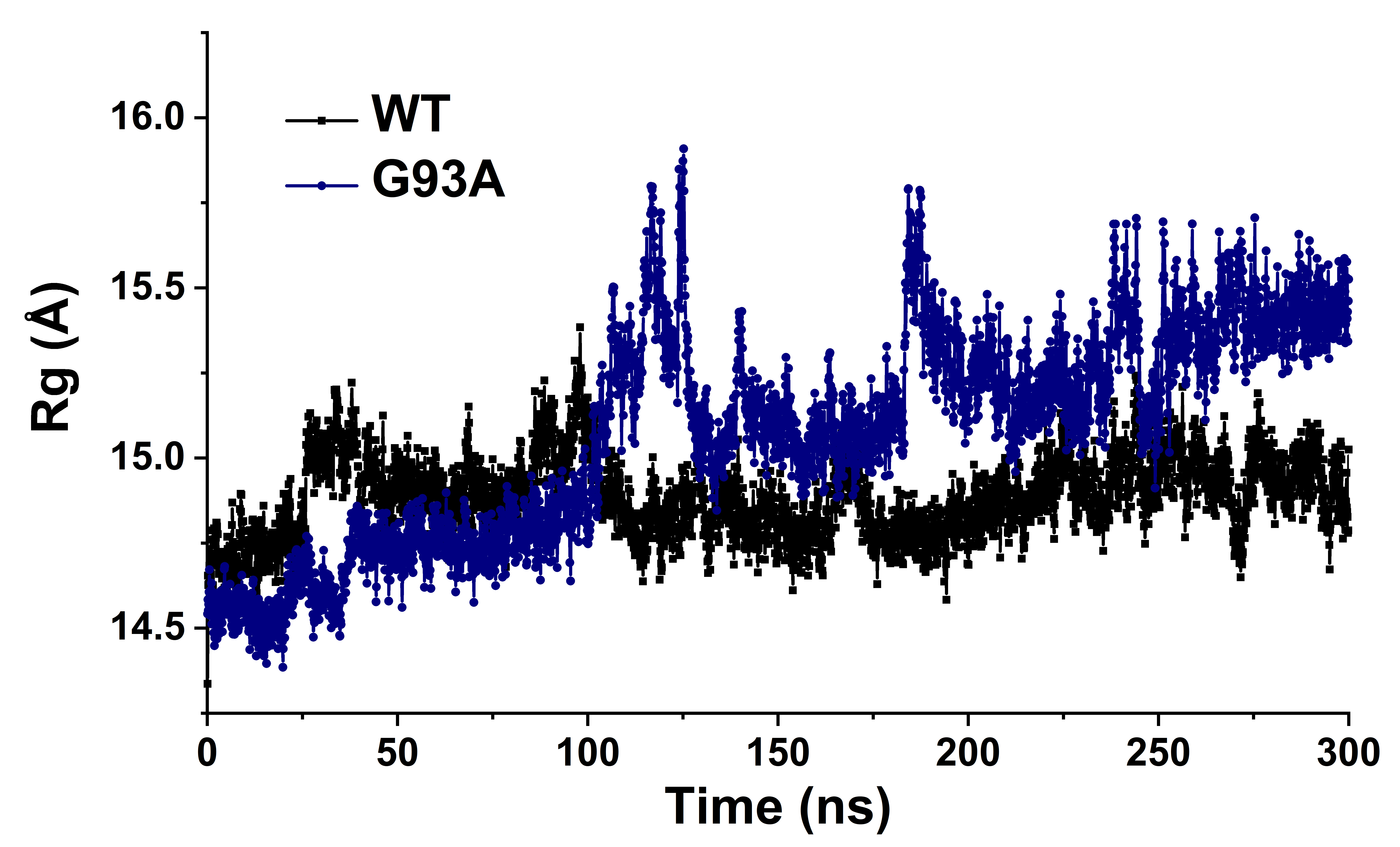

**Figure S8 |** Rg plot of **a**, WT-SOD1 and **b**, G93A-SOD1 systems.

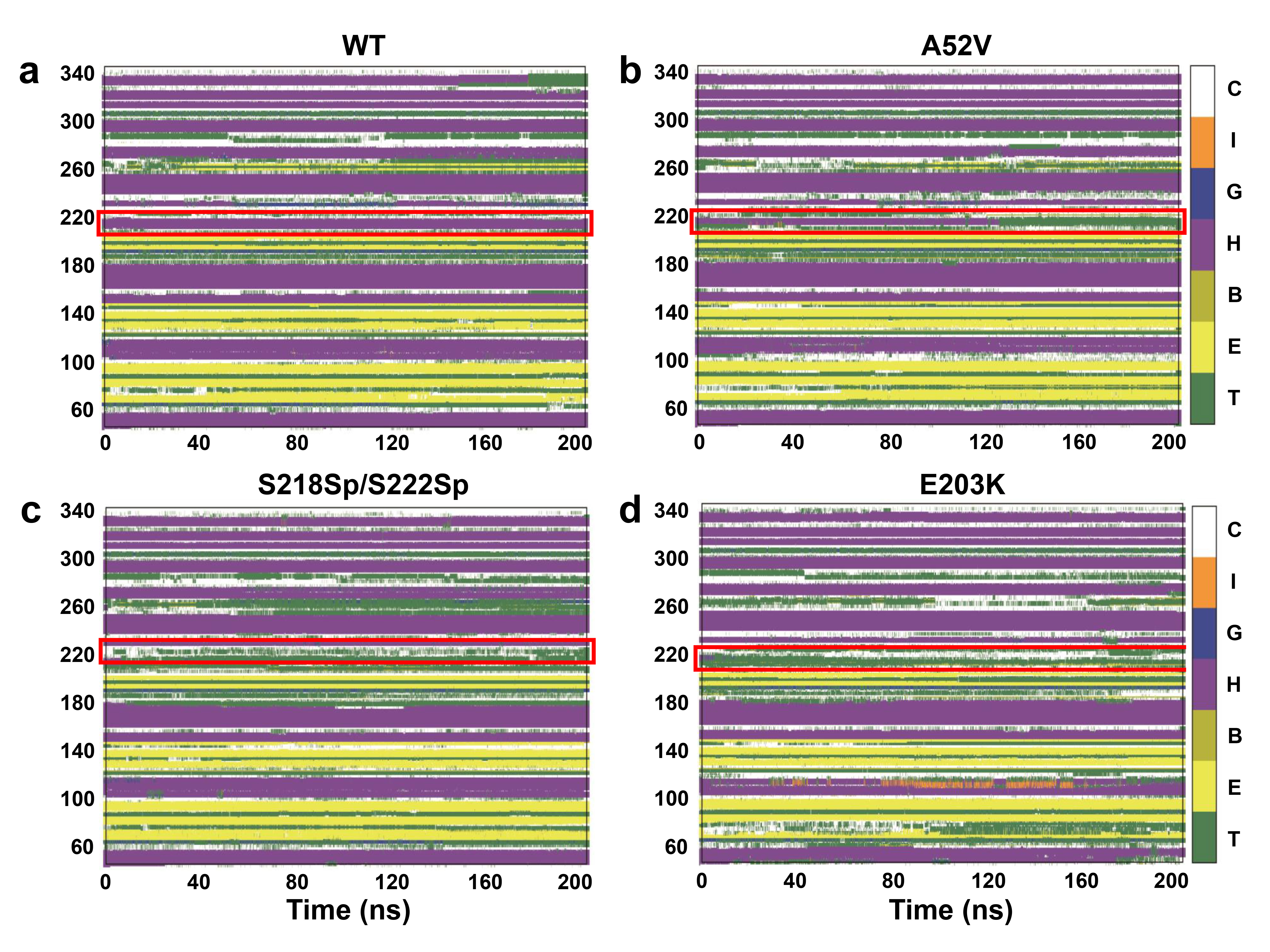

**Figure S9 |** Secondary structure analysis for **a**, WT, **b**, A52V, **c**, S218Sp/S222Sp, and **d**, E203K MEK1.

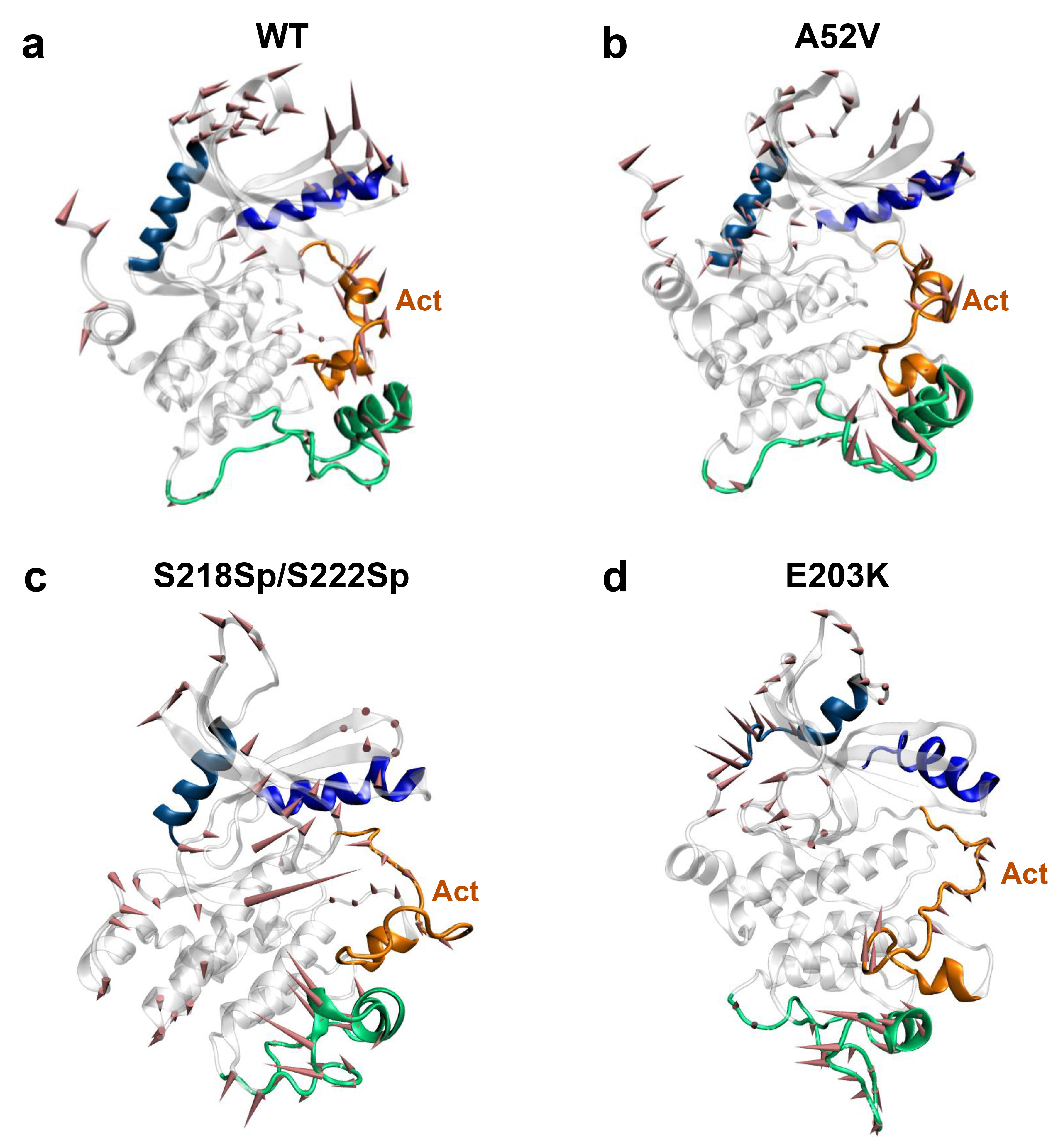

**Figure S10 |** Principal component analysis (PCA) for **a**, WT, **b**, A52V, **c**, S218Sp/S222Sp, and **d**, E203K MEK1. The direction shown in the cartoon denotes the motion mode of protein.

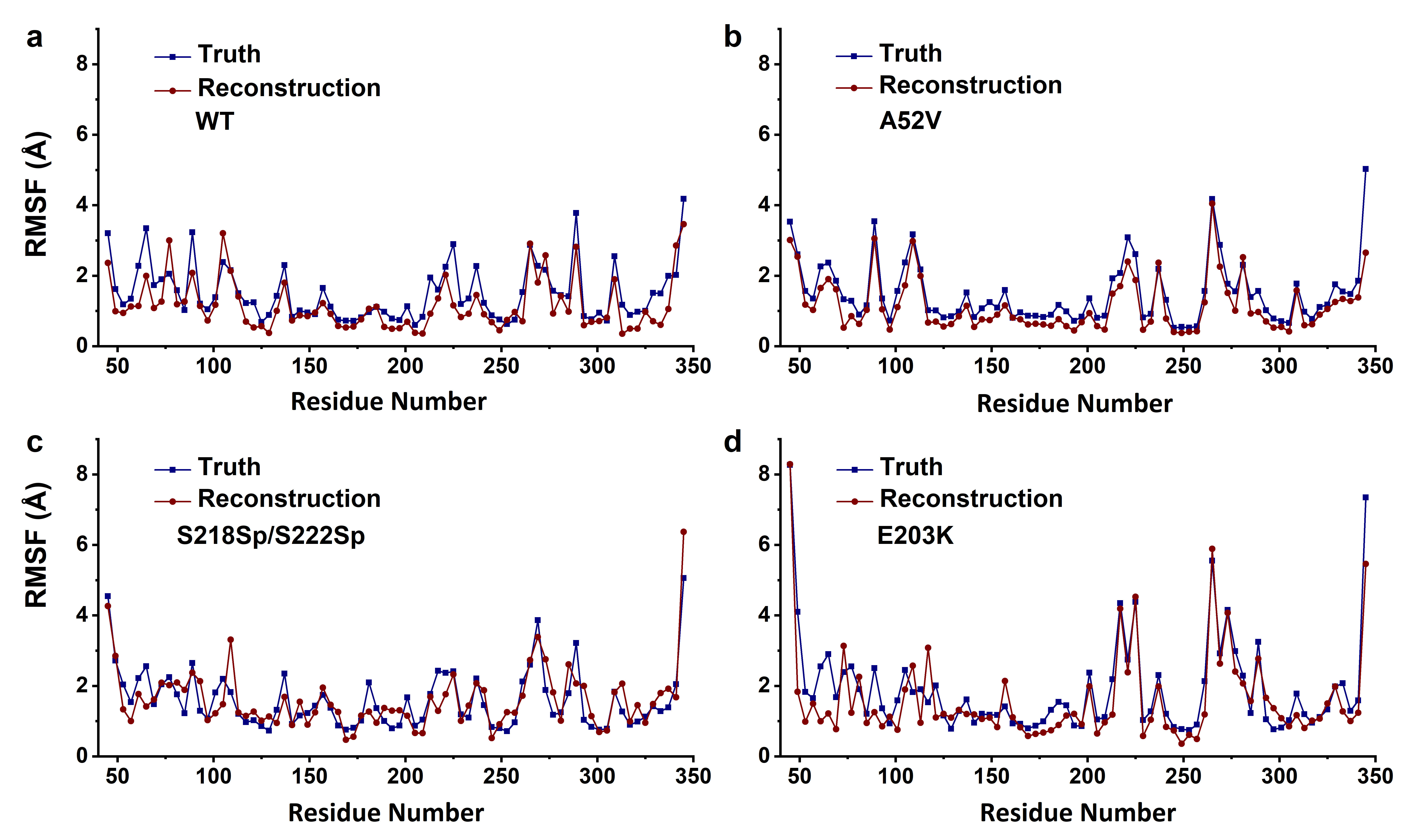

**Figure S11 |** Comparison of RMSF values between truth and reconstruction of trajectories for **a**, WT, **b**, A52V, **c**, S218Sp/S222Sp, and **d**, E203K MEK1.

**Table S1** **|** Mean squared error (MSE) in reconstructing trajectories with three interacting systems.

| **Pin1** | | **SOD1** | | **MEK1** | | | |
| --- | --- | --- | --- | --- | --- | --- | --- |
| Apo | 4.32e-3 | WT | 6.87e-3 | WT | 2.81e-3 | E203K | 7.15e-3 |
| FFpSPR-Pin1 | 4.10e-3 | G93A | 5.63e-3 | A52V | 2.71e-3 | S218Sp/S222Sp | 6.79e-3 |
| FFpSPR-Pin1 (I28A) | 4.02e-3 |  |  |  |  |  |  |

**Table S2 |** The shortest pathways from residues in WW domain to residues in catalytic loop for FFpSPR-Pin1, and from residues in WW domain to K97 in α1-helix for Apo-Pin1 and FFpSPR-Pin1 (I28A).

| **FFpSPR-Pin1** | **Apo-Pin1** |
| --- | --- |
| L7>Y23>M15>R17>R21>**K13>T29>Q131**>R69  P9>S41>P37>M15>R17>R21>**K13>T29>Q131**>R69  W11>M15>R17>R21>**K13>T29>Q131**>R69  **K13>T29>Q131**>R69  M15>R17>R21>**K13>T29>Q131**>R69  R17>R21>**K13>T29>Q131**>R69  S19>**K13>T29>Q131**>R69  R21>**K13>T29>Q131**>R69  Y23>M15>R17>R21>**K13>T29>Q131**>R69  F25>R21>**K13>T29>Q131**>R69  H27>M15>R17>R21>**K13>T29>Q131**>R69  **T29>Q131**>R69  A31>M15>R17>R21>**K13>T29>Q131**>R69  Q33>M15>R17>R21>**K13>T29>Q131**>R69  E35>M15>R17>R21>**K13>T29>Q131**>R69  P37>M15>R17>R21>**K13>T29>Q131**>R69  G39>S43>S41>P37>M15>R17>R21>**K13>T29>Q131**>R69  S41>P37>M15>R17>R21>**K13>T29>Q131**>R69  S43>S41>P37>M15>R17>R21>**K13>T29>Q131**>R69  L7>Y23>M15>R17>R21>**K13>P9>P133**>S67  **P9>P133**>S67  W11>M15>R17>R21>**K13>P9>P133**>S67  **K13>P9>P133**>S67  M15>R17>R21>**K13>P9>P133**>S67  R17>R21>**K13>P9>P133**>S67  S19>**K13>P9>P133**>S67  R21>**K13>P9>P133**>S67  Y23>M15>R17>R21>**K13>P9>P133**>S67  F25>R21>**K13>P9>P133**>S67  H27>M15>R17>R21>**K13>P9>P133**>S67  T29>Q131>S67  A31>M15>R17>R21>**K13>P9>P133**>S67  Q33>M15>R17>R21>**K13>P9>P133**>S67  E35>M15>R17>R21>**K13>P9>P133**>S67  P37>M15>R17>R21>**K13>P9>P133**>S67  G39>S43>S41>P37>M15>R17>R21>**K13>P9>P133**>S67  S41>P37>M15>R17>R21>**K13>P9>P133**>S67  S43>S41>P37>M15>R17>R21>**K13>P9>P133**>S67  L7>Y23>M15>**E35>K97>S105>C113**>K77  W11>M15>**E35>K97>S105>C113**>K77  K13>R21>M15>**E35>K97>S105>C113**>K77  M15>**E35>K97>S105>C113**>K77  R17>R21>M15>**E35>K97>S105>C113**>K77  S19>R21>M15>**E35>K97>S105>C113**>K77  R21**>**M15>**E35>K97>S105>C113**>K77  Y23>M15>**E35>K97>S105>C113**>K77  F25>R21>M15>**E35>K97>S105>C113**>K77  Q33>M15>**E35>K97>S105>C113**>K77  **E35>K97>S105>C113**>K77  P37>M15>**E35>K97>S105>C113**>K77  S41>P37>M15>**E35>K97>S105>C113**>K77 | **L7>E35>**K97  P9>**L7>E35**>K97  W11>**L7>E35**>K97  K13>R17>Y23>S19>**R21>E35**>K97  M15>**R21>E35**>K97  R17>Y23>S19>**R21>E35**>K97  S19>**R21>E35**>K97  **R21>E35**>K97  Y23>S19>**R21>E35**>K97  F25>**R21>E35**>K97  H27>**L7>E35**>K97  T29>**L7>E35**>K97  A31>**L7>E35**>K97  Q33>M15>**R21>E35**>K97  P37>M15>**R21>E35**>K97  G39>**L7>E35**>K97  S41>G39>**L7>E35**>K97  S43>G39>**L7>E35**>K97 |
|  | **FFpSPR-Pin1(I28A)** |
|  | L7>K13>M15>**R21>E35**>K97  P9>K13>M15>**R21>E35**>K97  W11>M15>**R21>E35**>K97  K13>M15>**R21>E35**>K97  M15>**R21>E35**>K97  R17>M15>**R21>E35**>K97  S19>**R21>E35**>K97  **R21>E35**>K97  Y23>**R21>E35**>K97  F25>M15>**R21>E35**>K97  H27>M15>**R21>E35**>K97  T29>K13>M15>**R21>E35**>K97  A31>M15>**R21>E35**>K97  Q33>M15>**R21>E35**>K97  **E35**>K97  P37>M15>**R21>E35**>K97  G39>F25>M15>**R21>E35**>K97  S41>P37>M15>**R21>E35**>K97  S43>G39>F25>M15>**R21>E35**>K97 |

**Table S3 |** The shortest pathways from G93/A93 to residues in EL loop for WT and G93A SOD1.

| **WT** | **G93A** |
| --- | --- |
| G93>**P13>K9>N53>A55>G61>T137>G73**>A123  G93>**P13>K9>N53>A55>G61>T137>G73**>D125  G93>**P13>K9>N53>A55>G61>T137**>E133>G127  G93>**P13>K9>N53>A55>G61>T137**>G129  G93>**P13>K9>N53>A55>G61>T137>G73**>T135>N131  G93>**P13>K9>N53>A55>G61>T137**>E133  G93>**P13>K9>N53>A55>G61>T137>G73**>T135  G93>**P13>K9>N53>A55>G61>T137**  G93>**P13>K9>N53>A55>G61>T137**>E133>N139 | A93>G41>A123  A93>**P13>D11>R143>T137>E133**>T135>D125  A93>**P13>D11>R143>T137>E133**>N131>G127  A93>**P13>D11>R143>T137>E133**>N131>G129  A93>**P13>D11>R143>T137>E133**>N131  A93>**P13>D11>R143>T137>E133**  A93>**P13>D11>R143>T137>E133**>T135  A93>**P13>D11>R143>T137**  A93>**P13>D11>R143>T137>E133**>N131>G129>N139  A93>**P13>D11>R143>T137>E133**>G141 |

**Table S4** **|** Probability of hydrogen bond formation in the trajectories of WT and G93A SOD1.

| **Donor** | **Acceptor** | **WT (%)** | **G93A (%)** |
| --- | --- | --- | --- |
| L38: N | G93/A93: O | 11.52 | 5.09 |
| H43: NE2 | T39: O | 23.54 | 28.05 |
| G44: N | H120: O | 20.57 | 17.35 |
| K122: NZ | A140: O | 6.82 | 1.38 |
| R115: NH1 | E49: O | 23.57 | 5.79 |
| Q22: NE2 | S105: OG | 30.06 | 15.58 |
| V47: N | G82: O | 36.67 | 24.64 |
| T116: OG1 | F50: O | 67.29 | 45.84 |
| D124: N | N86: OD1 | 26.18 | 17.85 |
| R79: NH2 | D101: OD1 | 24.65 | 20.17 |

**Table S5 |** The shortest pathways from N221 in the activation segment to residues in αA-helix and proline-rich loop in the S218Sp/S222Sp MEK1, and from R201 (near to E203K) to αC-helix and proline-rich loop in the E203K MEK1.

| **S218Sp/S222Sp** | **E203K** |
| --- | --- |
| N221>**G225>G77>E73>E69>P89>K57**>R49>Q45  N221>**G225>G77>E73>E69>P89>K57**>R49  N221>G225>G77>E73>E69>P89>V85>V93>F53  N221>**G225>G77>E73>E69>P89>K57**  N221>**D277**>S265>Y261  N221>**D277**>S265  N221>F273>S269  N221>F273  N221>**D277**  N221>**D277**>N281  N221>**D277**>S265>P285  N221>**D277**>S265>P285>S289  N221>**D277**>S265>P285>S289>S293 | R201>**Q45>R49>K57>V117**>N109>P105  R201>**Q45>R49>K57>V117**>N109  R201>**Q45>R49>K57>V117**>R113  R201>**Q45>R49>K57>V117**  R201>Q45>R49>C121  R201>**Q45>R49>K57>V117>G213>D217>G237>F273**>S265>Y261  R201>**Q45>R49>K57>V117>G213>D217>G237>F273**>S265  R201>**Q45>R49>K57>V117>G213>D217>G237>F273**>S269  R201>**Q45>R49>K57>V117>G213>D217>G237>F273**  R201>**Q45>R49>K57>V117>G213>D217>G237**>D277  R201>**Q45>R49>K57>V117>G213>D217>G237**>N281  R201>F333>E329>S293 |

**Supplementary Video 1** **|** Comparison of truth and reconstruction for apo Pin1, FFpSPR bound Pin1, and FFpSPR bound Pin1 (I28A).

**Supplementary Video 2** **|** Comparison of truth and reconstruction for WT-SOD1 and G93A-SOD1.

**Supplementary Video 3** **|** Comparison of truth and reconstruction for WT, A52V, S218Sp/S222Sp, and E203K MEK1.
